# The evolution and mechanistic versatility of the bacterial NADH dehydrogenases type II

**DOI:** 10.1101/2025.10.14.682086

**Authors:** Diego Masone, Lianne van Es, Guang Yang, Marco W. Fraaije, María Laura Mascotti

**Affiliations:** IHEM, CONICET-UNCuyo, M5502JMA Mendoza, Argentina; Molecular Enzymology group, University of Groningen, 9747AG Groningen, The Netherlands; School of Food Science and Pharmaceutical Engineering, Nanjing Normal University, 2 Xuelin Road, Nanjing, China

**Keywords:** NADH dehydrogenase type II, ancestral sequence reconstruction, evolution, charge transfer complex, Firmicutes

## Abstract

Type II NADH dehydrogenases (NDH-2s) are accessory enzymes of the bacterial electron transport chain (ETC). While they functionally overlap with Complex I, their main role is not proton translocation but maintaining the intracellular NADH/NAD^+^ balance. Although often non-essential, NDH-2 become crucial in species lacking complex I, serving as the primary electron entry point into the ETC. Their virtual absence in mammals makes these enzymes attractive targets for antimicrobial drug development and mitochondrial functional restoration. NDH-2s catalyze electron transfer from NADH to quinones, yet two distinct catalytic mechanisms have been described for members of the family: a classical ping-pong mechanism and an atypical ternary mechanism involving the formation of a charge transfer complex (CTC). The molecular basis of these mechanisms remains unclear. Also, their occurrence among NDH-2s from different bacterial lineages in unknown. Here we combined molecular phylogenetics, ancestral sequence reconstruction, expression and biochemical characterization of ancestral and modern enzymes and, molecular dynamics simulations to explore the mechanistic versatility of NDH-2s across Bacteria. Our results show the atypical ternary mechanism is restricted to the Firmicutes (Bacillota) lineage and it is defined by the presence of a single substitution located at the bottom of the active site. This work provides an evolutionary framework for understanding NDH-2 mechanistic versatility. Besides, it establishes a basis for drug discovery targeting pathogenic strains and opens avenues to develop innovative strategies to complement dysfunctional mitochondria.

## Introduction

Bacteria thrive in a wide range of ecological niches, largely due to their respiratory flexibility. This plasticity is enabled by branched electron transport chains (ETCs), which allow the use of alternative electron donors and acceptors depending on environmental and metabolic conditions [1]. Type II NADH dehydrogenases (NDH-2s) —also known as NADH quinone oxidoreductases— are accessory components of the ETC [2]. These enzymes transfer reducing equivalents from cytoplasmic NADH to the membrane quinone pool, functionally overlapping with complex I. However, unlike complex I, NDH-2s do not translocate protons across the membrane thus not contributing to the proton motive force. Their role is shifted towards balancing the intracellular balance of NADH/NAD^+^ [3]. Although generally non-essential, NDH-2 become crucial in organisms that lack complex I —such as the pathogens *Plasmodium falciparum, Mycobacterium leprae* and *Staphylococcus aureus*— serving as the primary entry point for electrons into the ETC. Importantly, the virtual absence of NDH-2 in mammals makes these enzymes attractive targets for antimicrobial drug development [4].

NDH-2s belong to the two dinucleotide binding domain (tDNBD) superfamily. They are monotopic proteins —monomeric or homodimeric— located in the plasma membrane of bacteria facing the cytoplasm, and use FAD as cofactor [2]. They display the typical arrangement of a flavin-binding domain and a nicotinamide binding-domain facing each other forming the active site at their interface. Besides, a C-terminal extension serves as anchor to the membrane and accommodates the substrate binding site —known as Q site— at the intersection between the anchoring point and the active site [5]. A few individual enzymes from the Bacteria domain have been biochemically characterized in different levels of detail. The two best studied are the *Caldalkalibacillus thermarum* NDH-2 (named here Ct-NDH2) and the *Staphylococcus aureus* NDH-2 (Sa-NDH2). Ct-NDH2 was the first enzyme in the family with an experimentally determined 3D structure [6] and later, its kinetic mechanism was thoroughly investigated [7]. In the same line, Sa-NDH2 has been structurally and mechanistically characterized [5, 8]. NDH-2s employ the coenzyme NADH as hydride donor [9], however the usage of NADPH *in vitro* has been demonstrated for Ct-NDH2 [7] and also speculated for uncharacterized sequences based on a single point mutation reported to change the hydride donor specificity [10]. Regarding the substrate, a variety of high and low redox potential quinones are accepted, making NDH-2s very diverse concerning their cellular roles [2]. Much work has been done on the discovery of novel inhibitors [11, 12]. However, these studies frequently emphasize the lack of structure-function rationale for candidate molecules due to the poor understanding of NDH-2s overall biochemistry. A reason for that is the scarcity of kinetic mechanisms characterized and some apparent contradictory results. The *Mycobacterium tuberculosis* NDH-2 (Mt-NDH2) shows a two-site ping-pong mechanism [13]. This is featured by a first step of flavin reduction via the hydride transfer from the NADH, followed by the release of the oxidized coenzyme NAD^+^. After that, the quinone substrate reaches the active site through a specific binding site and is subsequently reduced and released. This has been confirmed by fast-kinetic experiments as well as by the structural determination of two independent binding sites for the electron donor and acceptor species [13, 14]. Unlike the *Mycobacterium enzyme*, Ct-NDH2 and Sa-NDH2 follow an atypical ternary mechanism [5, 7]. This is characterized by the permanence of the NAD^+^ molecule in the active site during the whole catalytic cycle, enabling the formation of a UV-Vis active species called the charge transfer complex (CTC). The CTC is formed due to a π□π interaction between the isoalloxazine ring from the reduced FAD and the nicotinamide moiety of NAD^+^ and has a broad absorption band in the range of 600-700 nm [15]. Upon binding of the quinone, the CTC disappears as a consequence of the flavin oxidation via the substrate reduction. The cycle completes with the simultaneous release of NAD^+^ and the product. Fast-kinetics and mutagenesis analysis support this mechanistic proposal [8]. Over recent years, much of the research has been devoted towards a unified mechanistic understanding of NDH2s. For example, it has been described that Ct-NDH2 is able to follow a ping-pong mechanism *in vitro* if NADPH is provided instead of NADH [7], suggesting that there is an inherent tendency to switch among both kinetic modes in the enzyme family. Finally, the yeast enzyme, named Ndi1, shows a ternary mechanism as well. However, in this case the CTC is formed upon addition of NADH only when the substrate ubiquinone is already present in the active site, suggesting that a different network of interactions is established [16, 17].

NDH-2s are widespread, found not only in most bacterial classes but also in fungi, protists, and archaea [3, 10, 18]. Evolutionary history has been inferred by non-probabilistic methods, such as Neighbor Joining, and the topology of the tree does not follow the species tree. This suggests that events of duplication and horizontal gene transfer might have been frequent [10]. Marreiros *et al* proposed a classification based on phylogenetic clustering, depicting four groups named A-D. Group A includes a small number of uncharacterized sequences from proteobacteria, cyanobacteria and green algae, while group B is formed solely by eukaryotic sequences. Group C is formed almost exclusively by bacterial sequences (actinobacteria and proteobacteria classes) that have not been experimentally characterized. *In silico* structural analyses revealed that this group features a pair of cysteine residues predicted to form a disulfide bridge that may play a role widening the NADH pocket. Group D is the most extensively populated including the majority of the sequences characterized to date, either from prokaryotic or eukaryotic origin [10]. Several features, such as the presence of specific fingerprints or canonical residues were mapped revealing no clear trend in the clustering. However, a particularly interesting clue is the presence of NDH-2 largely biased towards species lacking other membrane-bound NADH: quinone oxidoreductases (as complex I or Na^+^-NQR). Given the abundance of NDH-2 in bacterial genomes and the scarcity in archaeal ones, a bacterial origin for the enzyme family was speculated [10]. Even more, phylogenomic analyses suggest that NDH-2s were part of the proteome of the last bacterial common ancestor [19]. From this body of knowledge, there are two clear observations, first the protein phylogeny is patchy and complex, and second, no single biochemical feature dictates the clustering in the tree.

A conundrum rises then: does the mechanism relate to taxonomic distribution among the bacteria lineages? To solve this question and rationalize the extent of the mechanistic diversity, we perform here the evolutionary-guided exploration of the NDH-2 family biochemistry. Employing phylogenetics, ancestral sequence reconstruction and biochemical characterization of ancestral and extant enzymes, we discovered that the formation of the charge transfer complex is specifically restricted to the Firmicutes (bacillota) species. Aided by molecular dynamics (MD) simulations we identified the sequence determinants that govern the enzyme’s catalytic mechanism, providing insights into the biological causes of this variation. We demonstrate that evolutionary clustering serves as a proxy for the mechanistic phenotype in future inhibitors design efforts.

## Results

### Evolutionary history of type II NADH:quinone oxidoreductases

The distribution of NDH-2s in Bacteria was investigated by analyzing the Database of Clusters of Orthologous Genes (COGs, [20]) COG1252: NADH dehydrogenase, FAD-containing subunit. Overall, most species contain a single copy of this gene while only some harbor duplications. The orthologous group includes some remote homologs (~20-30% identity) such as dihydrolipoamide dehydrogenases (DLDH) and sulfide:quinone oxidoreductases (SQR). To confidently vet true NDH-2s, we selected the sequences showing the two canonical motifs (Rossmann) [10] and excluded those displaying the ones described for DLDHs and SQRs [21, 22]. A representative dataset was obtained and employed to perform evolutionary analyses. The taxonomic distribution shows sequences in most of the Terrabacteria clades, with low representation in CPR (Candidate phyla radiation) group. In Gracilicutes lineages the distribution is patchy, as no sequences were detected in Verrucomicrobiota or Elusimicrobiota. Also, in the basal lineage Fusobacteriota no sequences were detected, whereas some homologs were found among the DST (Deinococcota, Synergistota and Thermotogota) group (Table S1). The recovered distribution agrees with the report of Marreiros *et al* [10] as well as with the inference of Coleman *et al*, including NDH-2s as part of the ancestral proteome of LBCA [19].

The phylogeny obtained by maximum likelihood inference method (unconstrained, Fig. 1 & Fig. S1) shows two major clades each formed by Terrabacteria and Gracillicutes species. This suggests that a possible pathway for the evolution of this family would start from two ancient paralogs in LBCA involving further specific duplications and losses. While major lineages as Actinobacteria and Proteobacteria form various clades spread across tree, Bacteroidota and Firmicutes are always recovered monophyletic. Aiming to recapitulate the evolution of the function of NDH-2, from early microbial populations to modern strains, we performed ancestral sequence reconstruction and selected seven nodes for experimental characterization (Fig. S2a). The rationale was to initially sample the broadest sequence space to investigate which aspects of the enzyme’s biochemistry are ancestral and which are innovations. The selected nodes correspond to well supported clades either formed by pure or mixed bacterial classes and the probabilities of the reconstruction varied from 0.76 to 0.87 (Fig. S2b). To account for an alternative evolutionary scenario of pure vertical inheritance, we constructed a second phylogeny constraining the protein tree with the bacterial species tree (Fig. S2c) and performed ASR. The ancestors of Proteobacteria (Anc1c, 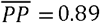), Acidobacteria (Anc2c, 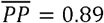), Firmicutes (Anc3c, 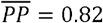), Bacteroidetes (Anc4c, 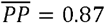) and Actinobacteria (Anc5c, 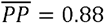) lineages were targeted (Fig. S2d).

**Fig. 1.**
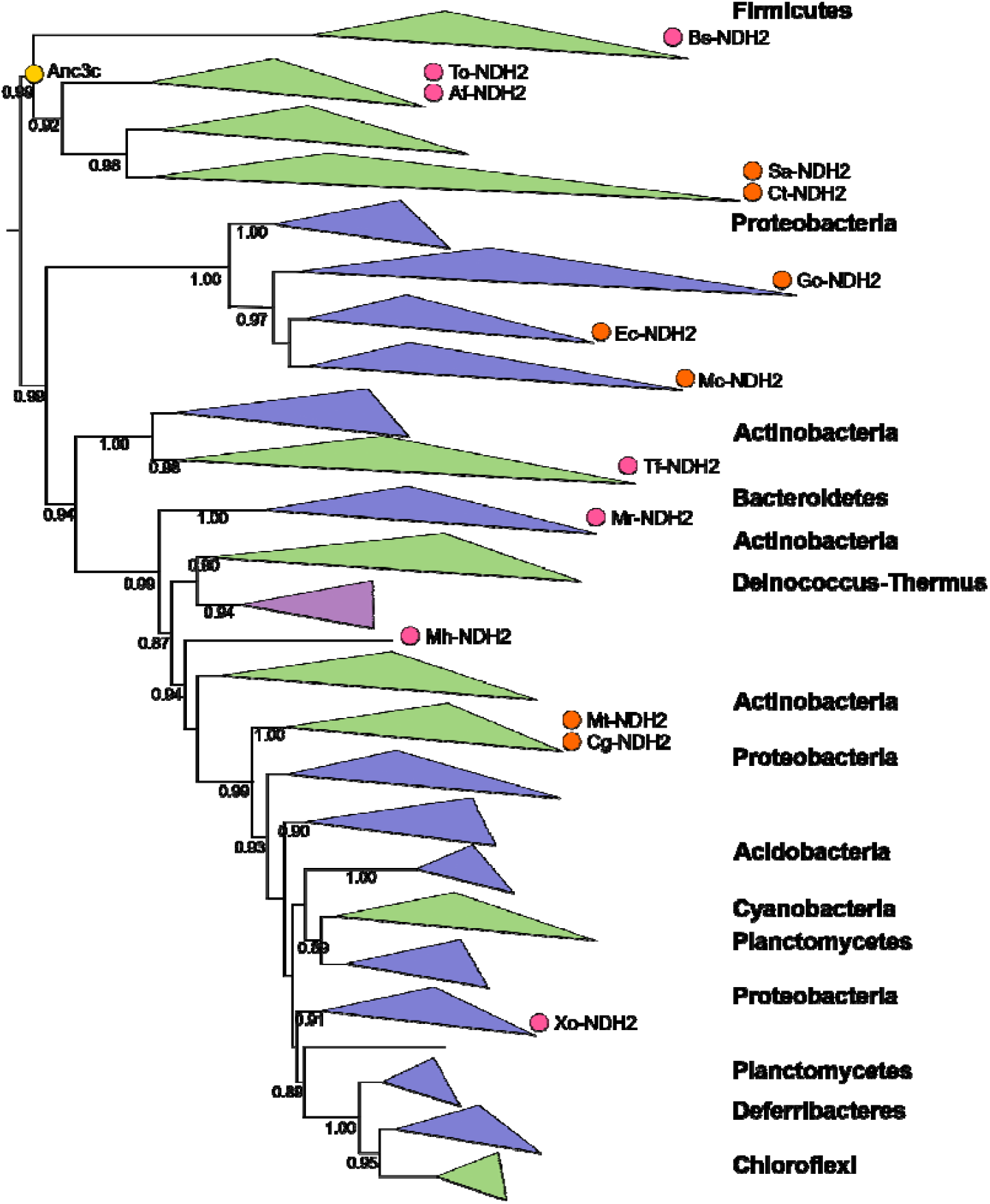
Phylogeny of the bacterial NDH-2 family. Tree was obtained in RaxML (MSA 493 sequences, 402 sites). It is shown as midpoint rooted for visual purposes. TBE values are shown at selected nodes. Color of clades indicate the bacterial clades: gracilicutes (blue), terrabacteria (green) and DST (violet). Names of the most representative classes for each clade are given on the right. Orange circles indicate extant enzymes previously characterized: Sa: *S. aureus*, Ct: *C. thermarum*, Go: *G. oxydans*, Ec: *E. coli*, Mc: *M. capsulatus*, Mt: *M. tuberculosis* and Cg: *C. glutamicum*. The yellow circle depicts Anc3c sequence. Pink circles show the novel enzymes targeted here Bs: *B. subtilis*, To: *T. oceani*, Af: *A. fermentans*, Tf: T. fusca, Mr: *M. ruestringensis*, Mh: *M. hydrothermalis* and Xo: *X. oceani*. Fully annotated phylogeny in FigS1.

### Hydride donor and substrate specificities are ancestral traits

Our collection of 12 ancestral NDH-2s representing early Bacteria evolution was cloned into pET and pBAD expression vectors. First, at small scale (50 mL), screening of various inducer concentrations and induction temperature/time regimes was performed. Out of 12, only six ancestral proteins were successfully expressed. As expected, these were found in the membrane fraction (Fig. S3). However, upon larger scale expression and purification two were not expressed in sufficient yields (Anc1c and Anc2c) and three were recovered as apo proteins, *i*.*e*. no FAD binding (Anc1u, 2u and 4u). Unfortunately, flavin reconstitution experiments were not successful [23]. Hence, only one ancestral enzyme, named Anc3c, corresponding to the Firmicutes clade, could be produced as holoenzyme with modest yields (4 mg/L culture) and good purity (Fig. 1 & Fig. S3). As control we expressed and purified three known NDH-2s: Ec-NDH2 from *Escherichia coli* [24], Mc-NDH2 from *Methylococcus capsulatus* [9] and Ct-NDH2 [7].

Steady-state kinetics indicate that Anc3c has high affinity for the endogenous substrate analogue menadione (*K*_*M*_ = 6.66 ± 2.4 µM) despite very low turnover compared to extant enzymes (*k*_*cat*_ = 0.86 ± 0.8 s^−1^). A similar behavior (high affinity, low turnover) was observed when other model quinones, such as benzoquinone and coenzyme Q1, were tested (Table 1, Fig. S4a). However, the *k*_*cat*_ was higher when coenzyme Q1 was employed as substrate, suggesting a more productive binding of this molecule at the active site.

**Table 1.**

|  | Anc3c | Mc-NDH2 | Ec-NDH2 | Ct-NDH2 |
| --- | --- | --- | --- | --- |
| *Steady state* |  |  |  |  |
| $k_{cat, \text{NADH}} [\text{s}^{-1}]$ | $0.77 \pm 0.07$ | $9.08 \pm 1.1$ | $43.6 \pm 4$ | $826.3 \pm 73.4$ |
| $k_{cat, \text{MD}} [\text{s}^{-1}]$ | $0.86 \pm 0.08$ | $3.9 \pm 0.4$ | $107.1 \pm 13.1$ | $409.1 \pm 36.7$ |
| $K_M, \text{NADH} [\mu\text{M}]$ | $58.8 \pm 13$ | $9.5 \pm 5.3$ | $1.7 \pm 1.09$ | $49.4 \pm 12.6$ |
| $K_M, \text{MD} [\mu\text{M}]$ | $6.7 \pm 2.4$ | $12.9 \pm 5.4$ | $15.6 \pm 8.5$ | $10.9 \pm 4.2$ |
| $k_{obs, \text{MD}} [\text{s}^{-1}]$ | $2.1 \pm 0.5$ | $10.1 \pm 0.004$ | $95.4 \pm 20.5$ | $425.5 \pm 31.4$ |
| $k_{obs, \text{coQ1}} [\text{s}^{-1}]$ | $4.6 \pm 0.5$ | $12.4 \pm 1.8$ | $267.5 \pm 183.6$ | $499.9 \pm 10.8$ |
| $k_{obs, \text{BQ}} [\text{s}^{-1}]$ | $1.28 \pm 0.2$ | $10.4 \pm 2.9$ | $33.4 \pm 0.2$ | $257 \pm 2^{(*)}$ |
| *Pre-steady state* |  |  |  |  |
| $k_{red} [\text{s}^{-1}]$ | $1.18 \pm 0.26$ | n.d | $18.6 \pm 2.3$ | $< 0.001$ |
| $k_{ox} [\text{s}^{-1}]$ | $21.1 \pm 0.9$ | n.d | $31.4 \pm 1.6$ | $> 231.3 \pm 36$ |
| $K_d, \text{NADH} [\mu\text{M}]$ | n.d | n.d | $> 12$ | n.d |
| $k_{unc} [\text{s}^{-1}]$ | n.d | n.d | n.d | $0.0123 \pm 0.0005$ |

MD: menadione, coQ1: coenzyme Q1, BQ: benzoquinone
$k_{obs, \text{MD}}$ , $k_{obs, \text{coQ1}}$ and $k_{obs, \text{BQ}}$ : are reported at 100 $\mu\text{M}$ substrate concentration and 0.1 $\mu\text{M}$ Anc3c, 10 nM Mc-NDH2 and 1nM Ct-NDH2 or Ec-NDH2.
(\*) at 50 $\mu\text{M}$ substrate concentration.

Since NDH-2s are widely spread across the Bacteria domain, and in some species lacking other membrane associated NADH:quinone oxidoreductases they are even the entry point to oxidative phosphorylation [10], we speculated that ancestral enzymes might display hydride donor promiscuity. The redox potential of Anc3c (□170.9 ± 0.3 mV) was found to be more positive than those of extant enzymes and even higher than free FAD (□219 mV). This can be seen as a sign of a preference to accept electrons, rather than to release them. Therefore, NADPH and F_420_H_2_ were tested as alternative electron donors. NADPH showed similar turnover at concentrations up to 50 µM (Fig. 2a), however kinetics could not be fitted to regular kinetic models (Fig. S4a). This indicates that in principle NADPH could deliver electrons to Anc3c, although not following a conventional mechanism. Anc3c was inactive when F_420_H_2_ was tested. These results show that the Firmicutes ancestral NDH-2 was a slow enzyme, though highly specific for quinone substrates and promiscuous for the nicotinamide nucleotide hydride donor.

**Fig. 2.**
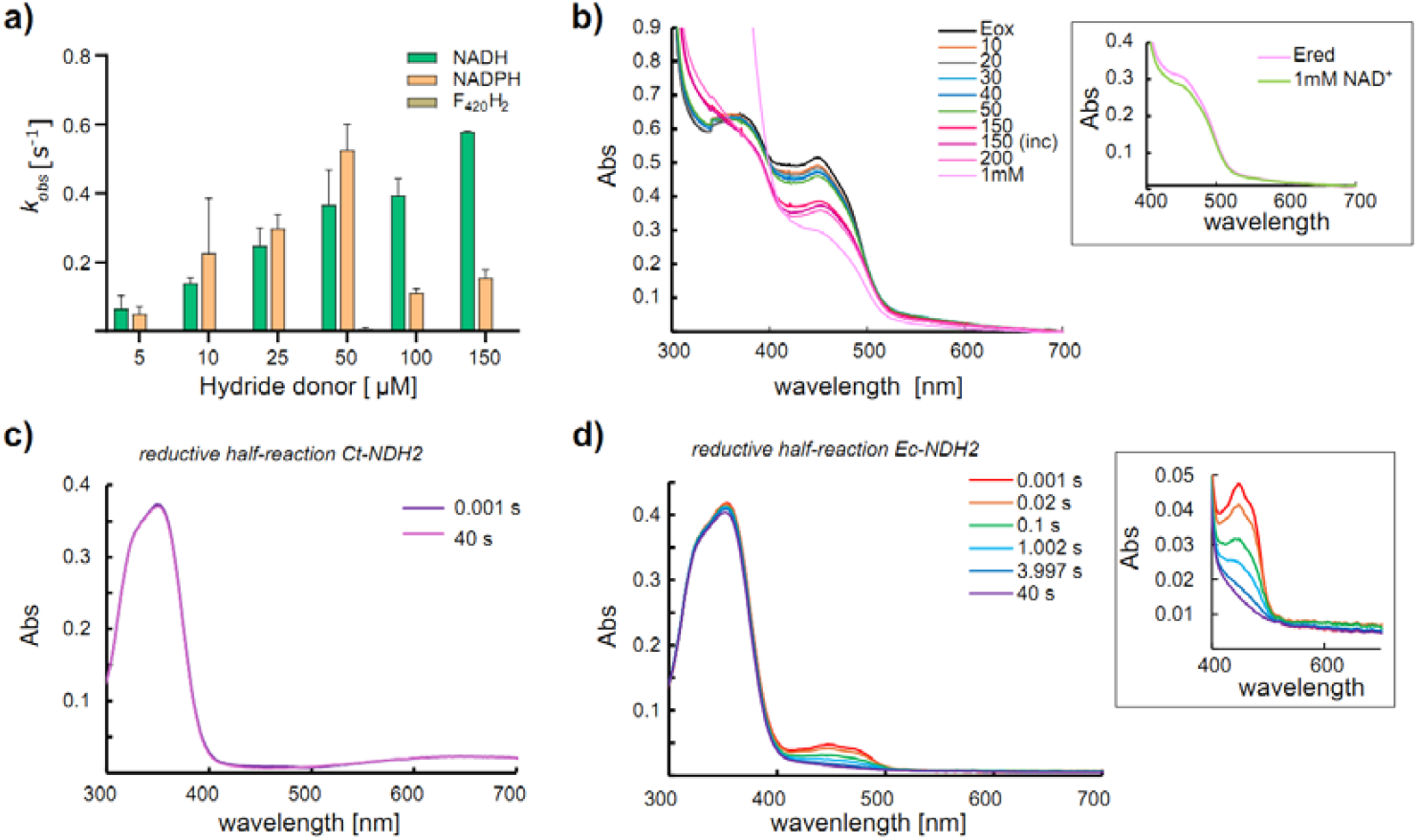
Biochemical characterization of Anc3c. **a)** Hydride donor preference of Anc3c. 0.75 µM of enzyme were used; **b)** NADH titration experiment of Anc3c. 44.25 µM of enzyme were used. The inset shows selected traces of the NAD^+^ titration at the high wavelength region where no absorption band was recorded; **c)** Selected traces of the reductive-half reaction of Ct-NDH2. The formation of the CTC is evidenced by a broad absorption band at 600-700 nm; **d)** Selected traces of the reductive-half reaction of Ec-NDH2. The inset shows a zoom of the high wavelength region where no absorption band was registered.

Kinetic features of the enzymes selected as controls showed a high degree of variation (Fig. S4b). Mc-NDH2 and Ec-NDH2 showed affinity values for menadione in the same order (~14 µM), however the *E. coli* enzyme was orders of magnitude faster (Table 1). Ct-NDH2 displayed the highest turnover values measured, reaching ~800 s^−1^ for NADH (*K*_*M*, NADH_= 49.4 ± 12.6 µM) and 400 s-1 (*K*_*M*, MD_= 10.9 ± 4.2 µM) for menadione. Reports indicate that enzymes showing an atypical ternary mechanism, display relatively high *k*_*cat*_ values and very similar µM values for NADH and its preferred substrate [7, 25].

### Extant enzymes showcase diverse catalytic mechanisms

The catalytic mechanism of NDH-2s has been debated in literature [13, 14]. Current evidence indicates that the Firmicutes Ct-NDH2 and Sa-NDH2 show an atypical ternary mechanism involving two sequential steps of NADH and substrate binding □hence atypical□ and the formation of the charge transfer complex (CTC) [7]. In contrast, the actinobacterial Mt-NDH2 shows a ping-pong mechanism [13] (Scheme 1). The reductive half-reaction analysis revealed that Anc3c is poorly reduced by NADH (~10%) in an apparent concentration independent manner, with a *k*_*red*_= 1.18 s^−1^. The absorbance spectrum formed upon reduction does not show a band compatible with the formation of the CTC. Titration with NADH in aerobic conditions, and absence of an electron acceptor, showed that nearly 30% of enzyme fraction could be reduced and confirmed that the CT complex is not formed (Fig. 2b). The oxidative half-reaction of this enzyme was analyzed upon anaerobic chemical reduction followed by mixing with the substrate menadione (Fig. S5a). The obtained oxidation rate (*k*_*ox*_= 21.1 s^−1^) suggests that this is not a rate limiting step. These features indicate that Anc3c does not follow a ternary mechanism involving a highly stable CTC. Unfortunately, this ancestor proved to be prone to aggregation, and this complicated further mechanistic characterization.

As only three bacterial NDH-2s have been mechanistically characterized —i.e.: Ct-NDH2, Sa-NDH2 and Mt-NDH2— we performed the catalytic cycle analysis of the *E. coli* enzyme (Ec-NDH2) in parallel with Ct-NDH2. In our hands the reduction rate exhibited by Ct-NDH2 was faster than the deadtime of the apparatus (< 0.001 s) and the formation of the CTC [Ct-NDH2·NAD^+^] was evidenced by a broad absorbance band at 600-700 nm (Fig. 2c). A key observation is the stability of the CTC towards O_2_ exposure, exhibiting oxidation of the flavin at *k*_*unc*_ = 0.012 s^−1^ while the CTC remains in place (Fig. S5b). The oxidative half-reaction showed the consumption of the [Ct-NDH2·NAD^+^] complex upon mixing with 25 µM of menadione at a *k*_*ox*_ > 230 s^−1^, indicating that this step likely determines the rate of catalysis (Fig. S5c). In contrast, the *E. coli* enzyme showed a significant slower reduction rate (*k*_*red*_ = 18.6 s^−1^), and no formation of the CTC (Fig. 2d). Like Anc3c, the enzyme was not fully reduced by an equimolar amount of NADH thus, to assess the oxidative half-reaction, it was chemically reduced. The oxidation of the reduced enzyme species, upon mixing with 50 µM of menadione, proceeded with a *k*_*ox*_ > 30 s^−1^, indicating that this step contributes partially to the turnover rate (Fig. S5d). Overall, these data suggest that Ec-NDH2 follows a ping-pong mechanism like the *M. tuberculosis* enzyme.

### The formation of the charge transfer complex is a feature of the Firmicutes clade

Is the mechanism of extant NDH-2s linked to their taxonomic distribution? To tackle this question, we selected seven enzymes across the NDH-2 phylogeny belonging to uncharacterized clades, for experimental characterization (Fig. 1). This set includes sequences from *Xanthomonas oryzae* (Xo-NDH2, gammaproteobacteria), *Muricauda ruestringensis* (Mr-NDH2, bacteroidetes), *Thermobifida fusca* (Tf-NDH2, actinobacteria), *Marinithermus hydrothermalis* (Mh-NDH2, deinococcus-thermus), *Anoxybacter fermentans* (Af-NDH2) and *Thermosediminibacter oceani* (To-NDH2, clostridia/firmicutes) and *Bacillus subtilis* (Bs-NDH2, firmicutes) (Table S2). Xo-NDH2 was not expressed while Mr-NDH2 was purified as apoprotein, the other five were expressed in good yields and successfully purified (Fig. S6).

A diagnostic of the atypical ternary mechanism occurrence is the spectrophotometric detection of the band in the 600-700 nm region corresponding to the CTC. Therefore, we conducted titration experiments with NADH. The actinobacterial enzyme Tf-NDH2 and the one from Deinoccocus-Thermus class Mh-NDH2 do not form the CTC upon exposure to increasing concentrations of NADH (Fig. 3, Fig. S7). Even more, Tf-NDH2 was not completely reduced by NADH, suggesting that either the oxidation by molecular oxygen is a fast process, or that the enzyme behaves as the one from *E. coli*. On the contrary, the three enzymes from the firmicutes clade Af-NDH2, To-NDH2 (both from clostridiales order) and Bs-NDH2, clearly showed the formation of the CTC when the NADH concentration was roughly double than enzyme (Fig. 3, Fig. S7). Considering previous reports and the data presented in this study, the ternary mechanism seems an exclusive feature of the firmicutes clade.

**Fig. 3.**
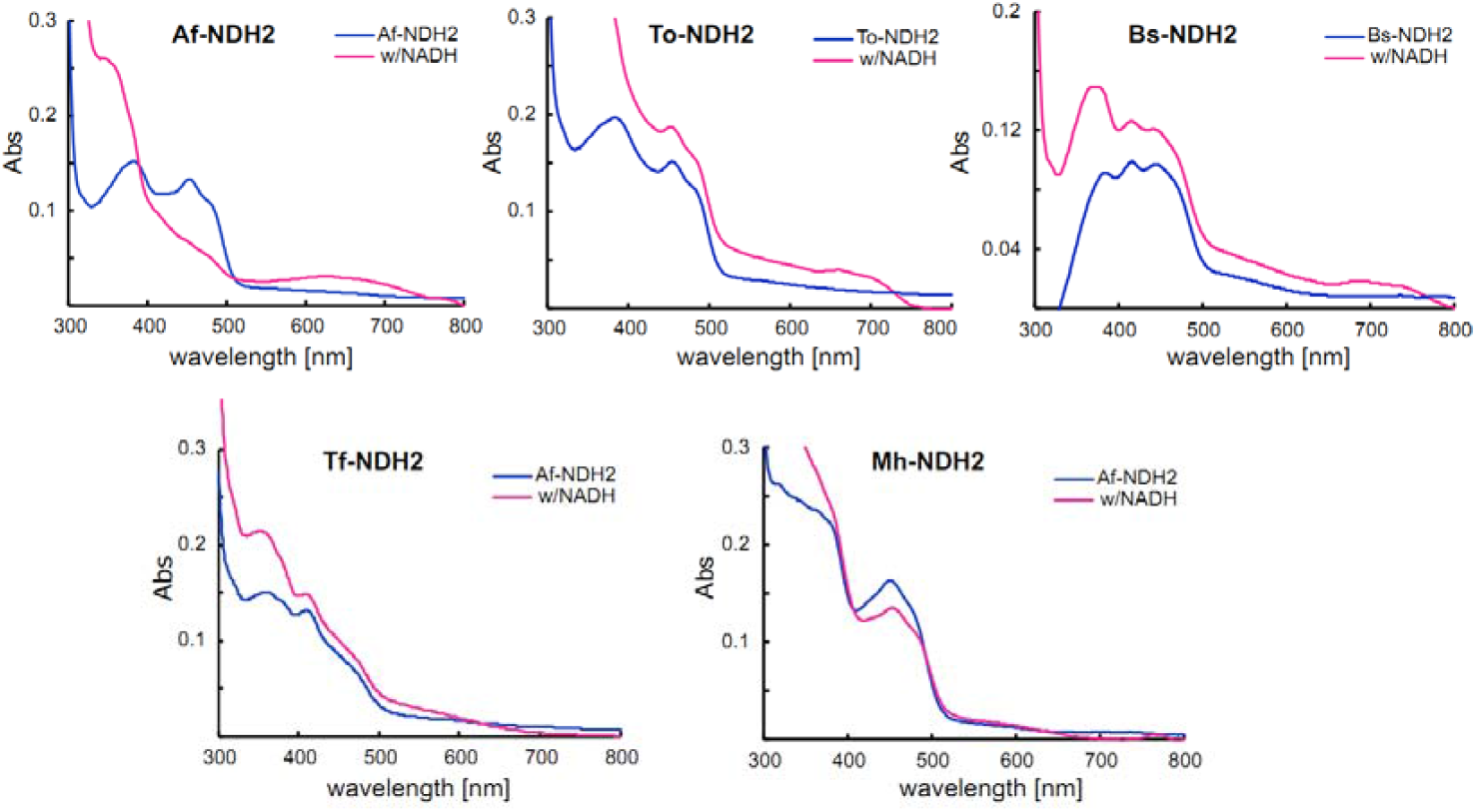
NADH titration experiments of newly characterized extant NDH-2. Top row shows enzymes from firmicutes class, forming the CTC upon mixing with 2x molar concentration of NADH. Bottom row shows enzymes from actinobacteria and DET groups, not forming the CTC. Enzyme concentration varied from 6-15 µM. Selected traces are shown (for the complete set please refer to Fig. S7).

### A single historical substitution enables CTC formation

To disclose the sequence determinants of CTC formation we performed a computational analysis involving structural analyses and atomistic molecular dynamics (MD) simulations employing either available structures or AlphaFold-predicted models, with FAD and NADH at the active sites. The Ct-NDH2 crystal structure (PDB: 5KMS) was used as well as its single mutant G164E (PDB: 5KMP) unable to form the CTC. On the other hand, we selected two enzymes preferring ping-pong mechanisms, the *M. tuberculosis* (Uniprot: P95160) and the *E. coli* (Uniprot: P00393). Anc3c was also included for which an AlphaFold model was generated. For all models, the two cofactors were extracted from Ct-NDH2 crystal structure to ensure no bias and further docked onto the active sites (Fig. S8a).

In flavoproteins, the formation of a CTC stable enough to be detected spectrophotometrically is the result of a π□π interaction among the isoalloxazine ring of the reduced flavin parallel stacked with the nicotinamide ring of NAD^+^ [15]. The G164E variant of *C. thermarum* NDH-2 provides an excellent tool in this analysis, as the mutation renders the protein unable to form it [7]. The glycine 164 residue is located at the second Rossmann motif defining the NADH binding site and it is conserved across the whole phylogeny. Upon structural inspection, the substitution of a glycine by a negatively charged amino acid generates a repulsion with the phosphate moiety of NAD(H), thus preventing the nicotinamide cofactor to reside long in the active site cavity. Hence, G164 is necessary but not sufficient to enable the CTC formation. To disclose the sequence determinants of the CT complex formation, we explored trajectories of 1 µs of unbiased MD simulations. The similar number of contacts among the protein and each of the cofactors, as well as the conserved mean value of the radius of gyration of NADH, show the health of our simulations (Fig. S8b). First, to confirm that we were able to capture the formation of the CTC, the angle (θ) between the planes formed by the aromatic rings of FAD and NAD was monitored across the entire simulation. Besides, the distance (d) between the nitrogen N8 from FAD molecule to the carbon C19 from NADH coenzyme was recorded (Fig. 4a). For extended aromatic systems, the slipped stacking is the most common type of geometry. This is featured by a mean distance of 3.8 Å among both centers and a 20° angle between the ring normal and the centroids vector [26]. As these values can suffer from deviations considering the contribution of other interactions to the system, especially in proteins, we defined the ranges 3.0-4.5 Å and 0-40 °, for distance and angle respectively. During the first half of the simulation (~0.5 µs) the CTC formation was confidently detected for Ct-NDH2 (Fig 4b) while the G164E variant showed a clear pattern disrupting the aromatic interaction, confirming the experimental phenotypes. The other proteins evaluated showed patterns coincident with their experimental spectroscopic behavior (Fig. S9a).

**Fig. 4.**
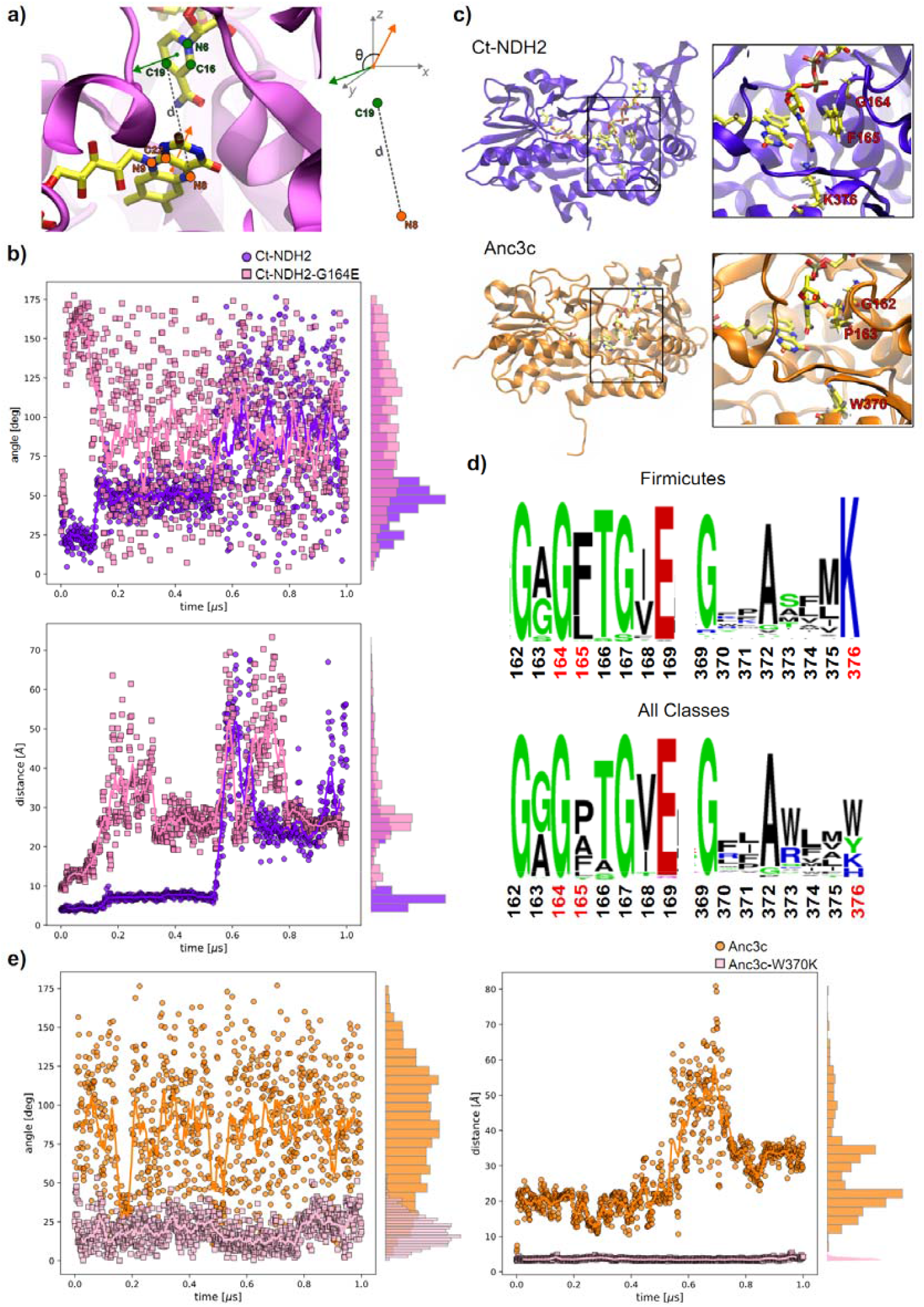
MD simulations and structural analysis. **a)** Scheme of angle/distance measurements. The image depicts the FAD plane defined by atoms N8, N9 and C22 (orange circles), the NADH plane defined by atoms N6, C16, C19 (green circles) and the normal vectors to each plane used to measure angle θ. Euclidean distance *d* among cofactors was defined between atoms N8 from FAD and C19 from NADH (dashed line); **b)** Top: angle measurements between the planes formed by the aromatic systems of FAD and NADH in Ct-NDH2 (5KMS, in violet) and Ct-NDH2-G164E (5KMP, in magenta) over 1 µs simulation time. Bottom: minimum distance measurements among FAD and NADH in Ct-NDH2 and Ct-NDH2-G164E over 1 µs simulation time; **c)** Representative structures of Ct-NDH2 (5KMS) and Anc3c (AlphaFold) with identified residues highlighted in yellow. The insets show a zoom in to the crucial substitutions. Proteins are shown in ribbons and FAD/NAD cofactors in bond representations with atoms colored as follows: C in yellow, O in red and N in blue; **d)** Sequence logo graphs of the Firmicutes sub dataset and the complete dataset. Residue numbering corresponds to Ct-NDH2. The identified substitutions are highlighted in red; **e)** Left: angle measurements between FAD and NADH planes in Anc3c model (in orange) and Anc3c-W370K model (in pink) over simulation time. Right: minimum distance measurements between FAD and NADH in Anc3c and Anc3c-W370K models over simulation time.

As the π□π interaction for the CTC formation depends on the protein microenvironment favoring the precise positioning of the two cofactors, we inspected the cavity surrounding both aromatic systems (~10 Å). Two crucial residues were spotted: F165 and K376 (numbering from Ct-NDH2 structure PDB: 5KMS, Fig. 4c). The first one —located in the second Rossmann fold— closely interacts with the nicotinamide moiety of NAD in Ct-NDH2 (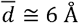 across 0.5 µs of the simulation), while in the G164E variant the larger average distance (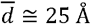 across 0.5 µs of the simulation) indicates no interaction along simulation time. For Mt-NDH2, a proline is found at the homologous site and although the distance was consistent across the entire simulation, it was larger, meaning no chances of aromatic interaction 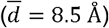.

In Ec-NDH2, the site is occupied by an alanine that after the 0.3 µs of the simulation acquires a constant 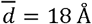. The phenylalanine substitution is not exclusive of the firmicutes clade □it is also found in some actinobacteria and gamma proteobacteria sequences. Therefore, although it seems to contribute to the stabilization of the complex, it is not essential (Fig. S9b). The conserved K376 is located at the C-terminal domain, pointing towards the interplanar section formed by the isoalloxazine ring and the nicotinamide moiety (towards the si-side of the flavin). In the enzyme microenvironment, this residue is predominantly protonated and thus may favorably interact with the flavin moiety. Simulations show that when Ct-NDH2 is impaired to form the ternary complex, K376 moves close enough to the flavin to prevent the π□π stabilization of the ternary complex (mean value 5.5 Å *vs* > 10 Å). The logo of the whole dataset shows that this lysine is strictly and exclusively conserved in the Firmicutes lineage (Fig. 4d). For other taxonomic groups aromatic residues are found at this site (Fig. S9c). Collectively, our results suggest that the simultaneous occurrence of at least G164 and K376 is sufficient to the CTC formation.

The fact that the ancestral NDH-2 from firmicutes does not form the CTC was puzzling. However, the structural analysis provided a clue. In Anc3c the first glycine is conserved, while the F165 is replaced by a proline (P163, PP= 1) and the K376 is substituted by a tryptophan (W370, PP=1). According to our analysis, the formation of the charge transfer complex would not be favored by this substitution network and instead catalysis would undergo via a ping-pong mechanism. To provide a rationale for this, we generated the single mutant Anc3c-W370K and performed MD simulations. From these simulations it became clear that the introduced substitution enabled the cofactors to interact with a mean distance of 3.8 Å and 20° angle (*vs* 20 Å and 100° in Anc3c) (Fig. 4e). Moreover, upon the single point substitution W370K, the proline 163 acquired a favorable 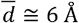 as observed in Ct-NDH2, compared to the wild-type variant that showed a two-phase behavior, with 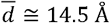 during the first half of the simulation, increasing by three-fold later. The lysine 370 shows a 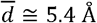 to the FAD molecule, suggesting a strong long-distance charge-charge interaction [27]. These results suggest that the historical substitution of an aromatic residue for a positively charged one, *i*.*e*.: lysine, in the Firmicutes lineage conferred the atypical ternary kinetic mechanism.

## Discussion

The mechanism of the NDH-2 enzyme family has long been a subject of debate. In this study, we address this issue from an evolutionary biochemistry perspective by exploring the sequence space within the Bacterial domain. By excluding eukaryotic sequences, we enhance the resolution of deep evolutionary events that have shaped the current distribution of NDH-2 enzymes. Our phylogenetic analysis reveals five distinct clades, broadly matching with the previously proposed groups [10]. However, through a more refined inference approach, we were able to further resolve those groups and ultimately identify a key feature underlying their divergence: differences in catalytic mechanism.

NDH-2’s main physiological role is to maintain the redox balance among NADH/NAD^+^ species. We evidenced that the specificity towards nicotinamide hydride donors is ancient. Interestingly, we were able to capture a level of promiscuity towards the phosphorylated version NADPH. This feature has not been completely erased in modern enzymes as in vitro they can effectively use NADPH as donor [7]. Conversely, NDH-2s show less specificity towards the identity of the quinone substrate, as they are able to reduce various menaquinone-derived structures [2]. Our results are in line with this observation. Interestingly, coenzyme Q1 —a version of the high potential quinones related to aerobic metabolism [28]— was well transformed by the ancestral NDH-2. This evidences an ancient metabolic versatility for the enzyme family.

Noticeable, the enzymes that overall show better catalytic performances are those following an atypical ternary mechanism. The detailed analysis of the reactivity with NADH provides an explanation for this. The experimental setup for measuring the oxidative half-reaction requires the preparation of the reduced enzyme which is typically achieved by reducing the enzyme by near equimolar amounts of the hydride donor in anaerobic conditions. Under these conditions, Ec-NDH2 shows only partial flavin reduction compared to Ct-NDH2, indicating a weak affinity of the enzyme for NADH (*K*_*d*_ >12 µM). A more pronounced situation is observed for Anc3c that is poorly reduced by NADH even in excess. In fact, the rate of reduction seems to be limiting the rate of catalysis for the enzymes operating with a ping-pong mechanism, while for the CTC-forming ones it is not the case [7]. For those, one of the most significant contributions to the overall reaction rate is the oxidative half-reaction, although the exact rate limiting step remains unknown. Interestingly, for other remote homologs —as the class B flavin monooxygenases— the rate limiting step is often the release of the oxidized nicotinamide cofactor occurring at the very end of the catalytic cycle [29], suggesting that such event could be a significant contribution to the catalytic rate of NDH-2.

The occurrence of the CTC formation, hence an atypical ternary mechanism of catalysis, seems to be restricted to the Firmicutes clade. Our structural analysis allowed to detect a clade-specific substitution of a lysine (K376, Ct-NDH2 numbering) at the C-terminal domain that in combination with the conserved glycine (G165) of the second Rossman-fold enables the formation of the complex. By using MD simulations, we were able to rationalize this mechanistic behavior. Previous bioinformatic analysis of the family shows that site 376 is among those with the highest cumulative covariance [30]. It was proposed to be involved in a proton conductive wire to the quinone pocket along with four carboxylic acids located at the nicotinamide binding domain. Our analysis shows that when this site is occupied by a long and positively charged residue, such as lysine, the distance to the N8/N6 atom of the flavin cofactor is such that it enables maintaining a charge-charge interaction that contributes to the stabilization of the complex. Also, we discovered that site 165, when occupied by an aromatic amino acid provides further stabilization to the CTC. Yet, this substitution is not crucial for the formation of the complex. The formation of the CTC complex, and thus catalysis via a ternary mechanism has been also demonstrated in the yeast homolog Ndi1 [17]. However, in Ndi1 the CTC is only formed upon the presence of both NADH and ubiquinone in the active site. In bacterial NDH-2s, the CTC is formed in a quinone substrate-independent manner upon reduction of the flavin by NADH. When substrate is present, the complex is consumed. Therefore, different stabilization mechanisms operate among the eukaryotic and bacterial homologs. Probably this is related to the two quinone binding sites observed in Ndi1 [16] which is unlike the bacterial enzymes. This evidence favors interpreting parallel evolution of the atypical ternary mechanism, rather than convergence. It has been speculated that the CTC formation would be an enzyme ‘adaptation’ to minimize the ROS production by the reduced flavin. Reduced flavins are known for their ability to produce ROS. This has been challenged by Blaza *et al* [7], due to the low magnitude of the NADH-driven rate of H_2_O_2_ production, likely insufficient to drive the evolution of the CTC. Our results also suggest that NDH-2 uncoupling is low regardless of the catalytic mechanism followed. This leaves the mechanistic question open. A clue is revealed by the turnover values. Firmicutes NDH-2s systematically show ~500-fold higher *k*_*cat*_ values for menadione derivatives than other NDH-2s, suggesting a highly efficient reduction of the quinone substrate. Therefore, CTC formation seems to enable a more efficient electron transfer by NDH-2s in maintaining cellular redox homeostasis as well as contributing to the electron transport chain. It could be speculated that, as the Firmicutes class is mostly composed of anoxygenic and facultative oxygenic species, these could benefit from a more efficient usage of cytoplasmatic hydride equivalents by their respiratory chains than other species.

Overall, our results provide a strong basis for pharmacological studies of NDH-2 as therapeutic targets in pathogenic strains. The phylogenetic clustering works as a proxy of the catalytic mechanism, hence knowing in advance such details of a target may enable smart design of novel inhibitors. Besides, our results open new avenues for strategies to complement malfunctioning mitochondria due to complex I failures.

## Materials and Methods

### Phylogenetic analysis

Bacterial NDH-2 sequences were retrieved from the COG1252 cluster (accession January 2022). The initial dataset included 992 sequences, among which remote homologs as dihydrolipoamide dehydrogenases (DLDH) and sulfide:quinone oxidoreductases (SQR) were found. After analysis of the presence of canonical fingerprints a representative and non-redundant dataset was obtained including 485 sequences. MAFFT v7 [31] was used to construct multiple sequence alignments (MSAs). MSAs were manually trimmed for single sequence extensions/insertions and gaps. RAxML v8.2.10 (HPC-PTHREADS module) [32] was employed to infer the maximum likelihood phylogeny with the fast-climbing algorithm (-f a), 450 bootstraps replicates, under gamma distribution and automated model selection. BOOSTER [33] was used to obtain the transfer bootstrap expectation (TBE) values as support. iTOL v7.2.1 [34] was employed to visualize and edit the tree. Bacterial species tree was obtained from TimeTree [35].

### Ancestral sequence reconstruction

Ancestral sequence reconstruction was performed using a reduced version of the phylogeny dataset (untrimmed MSA 362 sequences, 706 sites) and the tree topology was obtained as described in the previous section (unconstrained phylogeny). ASR was carried out as marginal reconstruction in pamlX v4.9a (codeml module) [36]. Sequences were analyzed using the LG substitution matrix, an empirical equilibrium amino acid frequency (model = 2), 16 gamma categories and re-estimation of α value. The posterior probability distribution at the targeted nodes was recorded. The length of the targeted ancestors was determined by parsimony. Sites were considered ambiguously reconstructed when PP < 0.8 and the second-best state displayed PP > 0.2. To assess the robustness of the reconstruction, the sequences of the alternative ancestors were also reconstructed [37]. ASR was also performed using a protein phylogeny constrained by the species tree.

### Chemicals and reagents

All NDH-2 encoding sequences were ordered as synthetic genes from Twist Bioscience. *E. coli* NEB 10β and BL21 competent cells, ligase, and restriction enzymes were purchased from New England Biolabs (NEB, USA). All other chemicals and enzymes were purchased from Sigma-Aldrich (Merck, USA) or Thermo Fisher (USA). Sequencing was carried out at Eurofins Genomics (Europe).

### Protein expression and purification

Ancestral and modern NDH-2s coding sequences were cloned employing the Golden Gate method. In house modified vectors, pBAD encoding a N-terminal 6xHis-SUMO tag (Amp^+^) and pET harboring an N-terminal 6xHis tag (Kan^+^), were used as recipient vectors. The cloning mixture contained 75 ng of recipient vector, ~40 ng of genes (2:1 molar ratio insert:vector), 15 U BsaI-HF, 15 U T4 DNA ligase, T4 ligase buffer and nuclease-free Milli-Q water up to a final volume of 20 µL. The mixtures underwent the following steps: i) 5 min at 37°C, ii) 10 min at 16°C (repeat 30 cycles both steps), iii) 10 min at 55°C and iv) 20 min at 65°C. 2 µL of the assembly mix were used to transform 50 µL of *E. coli* NEB 10-beta competent cells. The recovered cells were plated on LB-agar plates containing 50 mg/mL of appropriate antibiotic and incubated overnight at 37°C. Single colonies were picked to grow overnight in LB medium with antibiotic, and the plasmids were extracted using the QIAprep Spin Miniprep Kit (QIAGEN) and verified by Sanger sequencing. Glycerol stocks (25%) were stored at −80°C.

For expression of the pBAD constructs, 3 mL pre-cultures were prepared in LB medium with 50 mg/mL ampicillin overnight. To screen all constructs, 50 mL of LB medium containing antibiotic were inoculated with 0.5 ml of pre-culture and incubated at 37°C until reaching OD_600_= 0.4-0.6. L-arabinose was added to a final concentration of 0.02 % (w/v) to induce the cells and cultures were incubated overnight at 24°C, 200 rpm in a shaker for 20 h. After that, cells were harvested by centrifugation at 6000 rpm for 10 min. SDS-PAGE was used to confirm protein expression. For protein purification, constructs that showed expression were inoculated (2 mL of pre-culture) and induced in 1L flasks containing 200 mL of Terrific Broth medium containing 50 mg/mL ampicillin following the same conditions as before. After that, cells were harvested by centrifugation at 6000 rpm for 10 min. The pellets were resuspended in 100 mM KPi buffer, pH 7.5, containing 500 mM NaCl (Lysis Buffer) supplemented with 0.1 mM phenyl methyl sulfonyl fluoride and 3 ng/mL DNase. Cells were disrupted by sonication (70% amplitude, 5 sec on/ 5 sec off, for 10 min) and membrane fraction recovered by centrifugation at 11000 rpm for 60 min at 4°C (cell-free extract was discarded). Pellet was resuspended in Lysis Buffer supplemented with 0.5% (v/v) Triton X-100 and 100 µM FAD, and incubated overnight in the rotating wheel at 4°C. After this, solubilized membranes were obtained by centrifugation at 11000 rpm for 60 min at 4°C and recovering the soluble fraction. This was loaded into a column containing Ni-NTA resin (Thermo) equilibrated with Lysis Buffer. The non-bound proteins were washed out with 4 column volume (CV) of Lysis Buffer supplemented with 0.05% (v/v) Triton X-100, while the weakly bound proteins were removed with 4 CV of Lysis Buffer supplemented with 0.05% (v/v) Triton X-100 and 20 mM imidazole. Target proteins were eluted with Lysis Buffer supplemented with 0.05% (v/v) Triton X-100 and 500 mM imidazole. The eluted fractions with yellow color were pooled and loaded into a PD10 desalting column (Cytiva) to remove the imidazole and change the buffer to the Storage Buffer (50 mM KPi buffer, pH 7.5, 150 mM NaCl, 10 % glycerol and 0.05% Triton X-100). 30K filters were used to concentrate the proteins (Amicon). Protein concentration was determined by UV-Vis spectroscopy in a Jasco V-660 spectrophotometer based on the extinction coefficient of FAD (εFAD_440_ = 11.3 mM^−1^cm^−1^ at 450 nm). The purity was analyzed by SDS-PAGE and enzymes were frozen in small aliquots with liquid nitrogen and stored at –80°C.

For the pET constructs, 50 µL of *E. coli* BL21 were freshly transformed using 0.5-1 µL of plasmid and grown overnight in LB-agar plates at 37°C. Single colonies were picked the next day for pre-cultures preparation. Expression was conducted at 37°C for 6 hours, employing 1 mM IPTG.

### Thermal stability

ThemoFluor (39) and ThermoFAD (40) protocols were implemented in a BIORAD CFX Duet Real-Time PCR System. Enzyme concentrations ranged from 15-50 µM. The following protocol was used: from 25°C, 0.5/30 s to 90°C.

### Steady-state kinetics

Steady-state kinetic parameters were obtained by monitoring the consumption of NADH at 340 nm with a Jasco V-660 spectrophotometer. The observed rate constants were the initial rates calculated based on the molar extinction coefficient of NADH (εNADPH_340_= 6.22mM^−1^cm^−1^). The assays were done by mixing enzyme (final concentrations varying among enzymes from 1 nm to 1 µM) with varying substrate concentrations (0-100 µM) and NADH (0-300 µM) in Storage Buffer at 25°C. For kinetics on NADH, menadione was used as substrate and concentration was fixed at 100 µM. For kinetics towards quinone substrates, NADH concentration was 100 µM. The kinetic parameters were calculated with the fitting data using Michaelis-Menten equation in GraphPad Prism v 6.07. The uncoupling rates were determined in absence of substrates and increasing NADH concentration (0-150 µM). All measurements were done in triplicate, and rates are expressed as mean values ± standard deviation (SD).

### NADH/NAD^+^ titration

NADH/NAD^+^ affinity was determined by titration experiments. Stock solutions of enzymes (10-50 µM) and NAD+ (1-8 mM) were prepared. 1 µL of the NADH solution were added into NDH-2 solution to gradually reach a two times higher molar concentration than enzyme one, *i*.*e*.: from 0 to 200 µM. For NAD^+^ titration, final concentration reached 1 mM. The spectral changes were recorded in a Jasco V-660 or Shimadzu UV 1800 spectrophotometer every time after adding the NADH/NAD^+^ stock. The absorbance changes were plotted over wavelengths to detect the formation of the CTC complex in the 600-700 nm region. Experiments were done in duplicate.

### Redox potential measurement

The xanthine/xanthine oxidase method was employed (41). Anthraquinone-2,6-disulphonate (AQ26DS, E= □184mV) was found to be the best dye for Anc3c. The measurements were carried out using 8 µM of enzyme and 10 µM of dye in a Jasco V-660 spectrophotometer and spectra were collected for 30-50 cycles every 120 s. The absorbance changes at 480 nm were used for the calculations. Experiments were done in duplicate.

### Pre-steady-state kinetics

A SX20 stopped-flow spectrometer equipped with either a photodiode array (PDA) detector or a photomultiplier tube (PMT) module (Applied Photophysics, Surrey) was used. To generate anaerobic conditions, all reaction components were flushed with nitrogen for 10 min in sealed vials and supplemented with 5 mM glucose and 0.30 µM glucose oxidase (*Aspergillus niger* type VII).

The reductive half-reactions were first monitored using the PDA detector. 5-10 µM of enzyme were anaerobically reduced with excess of NADH (50-100 µM) and spectral changes were collected for 60 seconds. Every experiment represents the average of three independent measurements. Experiments were repeated three times. The reduction rates (*k*_*red*_) were obtained using the PMT detector at 450 nm. The absorbance change was recorded after mixing 10 µM enzyme and various NADH concentrations (0-200 µM) in anaerobic conditions. The observed rates (*k*_*obs*_) were obtained by fitting traces to exponential functions. The *k*_*red*_ and *K*_*d*_ values were estimated by fitting *k*_*obs*_ against NADH concentrations with Michaelis-Menten equation (eq1). All measurements were performed three times.

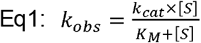

The oxidative half-reactions were monitored with the PDA detector. 5-10 µM of enzymes were fully reduced anaerobically with 1.5 equivalent NADH or sodium dithionite and then mixed with substrate. Spectral changes were collected for 60 seconds. Different concentrations of menadione substrate (25, 50 and 100 µM) were used. The *k*_*ox*_ values are the slopes of the curves obtained by plotting *k*_*obs*_ against substrate concentrations. To estimate the uncoupling rates, the same experiment was performed by mixing reduced enzymes with aerated buffer. The data were analyzed by Pro-Data Viewer v4.2.12, Pro-Kineticist v1.0.13 (Applied Photophysics) and GraphPad Prism v6.07 (Dotmatics). All experiments were repeated three times.

### Molecular dynamics

Structures were obtained from PDB and AlphaFold models from Uniprot. Anc3c model was obtained with AlphaFold2 [38]. Protein-ligand docking for FAD and NAD cofactors was conducted with the GRAMM-X webserver [39]. Atomistic simulations in the microsecond time scale were conducted with Gromacs [40] (version 2023). Systems were prepared using the CHARMM-GUI web server [41] (version 3.7), solvated in TIP3 explicit water and minimized/equilibrated following the CHARMM-GUI protocol for Gromacs [42, 43]. The pressure and the temperature were set to 1.0 bar and 298°K, and they were respectively controlled by a C-rescale barostat with a 5 ps time constant and a V-rescale thermostat with a 1 ps coupling constant. The Verlet cut-off scheme with Particle Mesh Ewald (PME) [44] was used with cut-offs set to 1.2 nm and a Force-switch vdw modifier at 1.0 nm. A 4th order LINCS algorithm was set for H-bond constraints [45].

Figures were prepared with Visual Molecular Dynamics (VMD) [46], Grace (GRaphing, Advanced Computation and Exploration of data) [47] and Inkscape [48].

#### Author contributions

All listed authors performed experiments and/or analyzed data. MLM conceived the idea with support of MWF. MLM performed molecular phylogenetics, ASR and set up the expression, purification and characterization of enzymes. LvE performed protein purification and kinetic experiments under MLM supervision. GY performed stopped flow experiments. DM performed docking and molecular dynamics simulations. MLM, DM and MWF analyzed the data. MLM wrote and edited the manuscript and prepared the figures with help from DM. DM, MWF and MLM revised the manuscript.

## Acknowledgements

This work was funded by: European Union’s Horizon 2020 Research and Innovation program under grant agreement No. 847675 COFUND project oLife (MLM); CONICET reallocation allowance (RESOL-2024-1665-APN-DIR#CONICET) to MLM and Grants from CONICET (PIP-0409CO) and ANPCyT (PICT2020-1897) to DM. Authors are grateful to students Tamara Otero and Valentina Alarcón who collaborated in the project, as well as Dr. Hein Wijma who provided initial input on structural analysis. Supercomputing time for this work was provided by CCAD-UNC (Centro de Computación de Alto Desempeño de la Universidad Nacional de Córdoba).

## Data availability

Constructs are available upon request to. Reconstructed ancestral sequences have been deposited in NCBI (accession codes: PX215251 (Anc1c), PX215252 (Anc2c), PX215253 (Anc3c), PX215254 (Anc4c), PX215255 (Anc5c), PX215256 (Anc1u), PX215257 (Anc2u), PX215258 (Anc3u), PX215259 (Anc4u), PX215260 (Anc5u), PX215261 (Anc6u) and PX215262 (Anc7u). Molecular dynamics files related to this work are freely available through GitHub at https://github.com/diegomasone/NDH2.

---

**Scheme 1.**
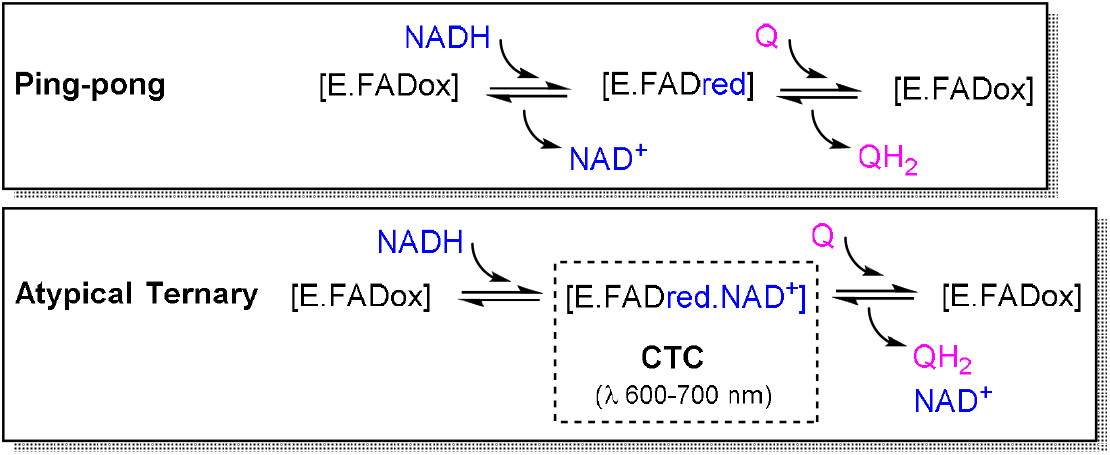
Simplified depiction of the two catalytic mechanisms described for NDH-2.

## References

1. Refojo, P.N., et al., The plethora of membrane respiratory chains in the phyla of life. Adv Microb Physiol, 2019. 74: p. 331–414.

2. Kerscher, S., et al., The three families of respiratory NADH dehydrogenases. Results Probl Cell Differ, 2008. 45: p. 185–222.

3. Melo, A.M., T.M. Bandeiras, and M. Teixeira, New insights into type II NAD(P)H:quinone oxidoreductases. Microbiol Mol Biol Rev, 2004. 68(4): p. 603–16.

4. Cook, G.M., et al., Energetics of pathogenic bacteria and opportunities for drug development. Adv Microb Physiol, 2014. 65: p. 1–62.

5. Sena, F.V., et al., Type-II NADH:quinone oxidoreductase from Staphylococcus aureus has two distinct binding sites and is rate limited by quinone reduction. Mol Microbiol, 2015. 98(2): p. 272–88.

6. Heikal, A., et al., Structure of the bacterial type II NADH dehydrogenase: a monotopic membrane protein with an essential role in energy generation. Mol Microbiol, 2014. 91(5): p. 950–64.

7. Blaza, J.N., et al., The mechanism of catalysis by type-II NADH:quinone oxidoreductases. Scientific Reports, 2017. 7(1): p. 40165.

8. Sousa, F.M., et al., The key role of glutamate 172 in the mechanism of type II NADH:quinone oxidoreductase of Staphylococcus aureus. Biochim Biophys Acta Bioenerg, 2017. 1858(10): p. 823–832.

9. Cook, S.A. and A.K. Shiemke, Evidence that a type-2 NADH:quinone oxidoreductase mediates electron transfer to particulate methane monooxygenase in methylococcus capsulatus. Arch Biochem Biophys, 2002. 398(1): p. 32–40.

10. Marreiros, B.C., et al., Type II NADH:quinone oxidoreductase family: phylogenetic distribution, structural diversity and evolutionary divergences. Environ Microbiol, 2016. 18(12): p. 4697–4709.

11. Mogi, T., et al., Identification of new inhibitors for alternative NADH dehydrogenase (NDH-II). FEMS Microbiol Lett, 2009. 291(2): p. 157–61.

12. Petri, J., et al., Structure of the NDH-2 - HQNO inhibited complex provides molecular insight into quinone-binding site inhibitors. Biochim Biophys Acta Bioenerg, 2018. 1859(7): p. 482–490.

13. Yano, T., et al., Mycobacterium tuberculosis Type II NADH-Menaquinone Oxidoreductase Catalyzes Electron Transfer through a Two-Site Ping-Pong Mechanism and Has Two Quinone-Binding Sites. Biochemistry, 2014. 53(7): p. 1179–1190.

14. Yano, T., et al., Steady-state kinetics and inhibitory action of antitubercular phenothiazines on mycobacterium tuberculosis type-II NADH-menaquinone oxidoreductase (NDH-2). J Biol Chem, 2006. 281(17): p. 11456–63.

15. Sakurai, T. and H. Hosoya, Charge-transfer complexes of nicotinamide-adenine dinucleotide analogues and flavin mononucleotide. Bibl Laeger, 1966. 112(3): p. 459–68.

16. Feng, Y., et al., Structural insight into the type-II mitochondrial NADH dehydrogenases. Nature, 2012. 491(7424): p. 478–482.

17. Yang, Y., et al., Reaction mechanism of single subunit NADH-ubiquinone oxidoreductase (Ndi1) from Saccharomyces cerevisiae: evidence for a ternary complex mechanism. J Biol Chem, 2011. 286(11): p. 9287–97.

18. Antos-Krzeminska, N. and W. Jarmuszkiewicz, Alternative Type II NAD(P)H Dehydrogenases in the Mitochondria of Protists and Fungi. Protist, 2019. 170(1): p. 21–37.

19. Coleman, G.A., et al., A rooted phylogeny resolves early bacterial evolution. Science, 2021. 372(6542): p. eabe0511.

20. Galperin, M.Y., et al., COG database update: focus on microbial diversity, model organisms, and widespread pathogens. Nucleic Acids Res, 2021. 49(D1): p. D274–d281.

21. Cherney, M.M., et al., Crystal structure of sulfide:quinone oxidoreductase from Acidithiobacillus ferrooxidans: insights into sulfidotrophic respiration and detoxification. J Mol Biol, 2010. 398(2): p. 292–305.

22. Mattevi, A., et al., Three-dimensional structure of lipoamide dehydrogenase from Pseudomonas fluorescens at 2.8 A resolution. Analysis of redox and thermostability properties. J Mol Biol, 1993. 230(4): p. 1200–15.

23. Hefti, M.H., J. Vervoort, and W.J. van Berkel, Deflavination and reconstitution of flavoproteins. Eur J Biochem, 2003. 270(21): p. 4227–42.

24. Björklöf, K., V. Zickermann, and M. Finel, Purification of the 45 kDa, membrane bound NADH dehydrogenase of Escherichia coli (NDH-2) and analysis of its interaction with ubiquinone analogues. FEBS Lett, 2000. 467(1): p. 105–10.

25. Schurig-Briccio, L.A., et al., Characterization of the type 2 NADH:menaquinone oxidoreductases from Staphylococcus aureus and the bactericidal action of phenothiazines. Biochim Biophys Acta, 2014. 1837(7): p. 954–63.

26. Janiak, C., A critical account on π–π stacking in metal complexes with aromatic nitrogen-containing ligands. Journal of the Chemical Society, Dalton Transactions, 2000(21): p. 3885–3896.

27. Zhou, H.X. and X. Pang, Electrostatic Interactions in Protein Structure, Folding, Binding, and Condensation. Chem Rev, 2018. 118(4): p. 1691–1741.

28. Franza, T. and P. Gaudu, Quinones: more than electron shuttles. Research in Microbiology, 2022. 173(6): p. 103953.

29. Torres Pazmiño, D.E., et al., Kinetic Mechanism of Phenylacetone Monooxygenase from Thermobifida fusca. Biochemistry, 2008. 47(13): p. 4082–4093.

30. Marreiros, B.C., et al., Structural and Functional insights into the catalytic mechanism of the Type II NADH:quinone oxidoreductase family. Scientific Reports, 2017. 7(1): p. 42303.

31. Katoh, K., J. Rozewicki, and K.D. Yamada, MAFFT online service: multiple sequence alignment, interactive sequence choice and visualization. Briefings in Bioinformatics, 2017. 20(4): p. 1160–1166.

32. Stamatakis, A., RAxML version 8: a tool for phylogenetic analysis and post-analysis of large phylogenies. Bioinformatics, 2014. 30(9): p. 1312–1313.

33. Lemoine, F., et al., Renewing Felsenstein’s phylogenetic bootstrap in the era of big data. Nature, 2018. 556(7702): p. 452–456.

34. Letunic, I. and P. Bork, Interactive Tree of Life (iTOL) v6: recent updates to the phylogenetic tree display and annotation tool. Nucleic Acids Research, 2024. 52(W1): p. W78–W82.

35. Kumar, S., et al., TimeTree: A Resource for Timelines, Timetrees, and Divergence Times. Molecular Biology and Evolution, 2017. 34(7): p. 1812–1819.

36. Yang, Z., PAML 4: Phylogenetic Analysis by Maximum Likelihood. Molecular Biology and Evolution, 2007. 24(8): p. 1586–1591.

37. Mascotti, M.L., Resurrecting Enzymes by Ancestral Sequence Reconstruction. Methods Mol Biol, 2022. 2397: p. 111–136.

38. Jumper, J., et al., Highly accurate protein structure prediction with AlphaFold. Nature, 2021. 596(7873): p. 583–589.

39. Tovchigrechko, A. and I.A. Vakser, GRAMM-X public web server for protein– protein docking. Nucleic Acids Research, 2006. 34(Suppl_2): p. W310–W314.

40. Abraham, M.J., et al., GROMACS: High performance molecular simulations through multi-level parallelism from laptops to supercomputers. SoftwareX, 2015. 1-2: p. 19–25.

41. Jo, S., et al., CHARMM-GUI: A web-based graphical user interface for CHARMM. Journal of Computational Chemistry, 2008. 29(11): p. 1859–1865.

42. Lee, J., et al., CHARMM-GUI Input Generator for NAMD, GROMACS, AMBER, OpenMM, and CHARMM/OpenMM Simulations Using the CHARMM36 Additive Force Field. Journal of Chemical Theory and Computation, 2016. 12(1): p. 405–413.

43. Qi, Y., et al., CHARMM-GUI Martini Maker for Coarse-Grained Simulations with the Martini Force Field. Journal of Chemical Theory and Computation, 2015. 11(9): p. 4486–4494.

44. Essmann, U., et al., A smooth particle mesh Ewald method. The Journal of Chemical Physics, 1995. 103(19): p. 8577–8593.

45. Hess, B., et al., LINCS: A linear constraint solver for molecular simulations. Journal of Computational Chemistry, 1997. 18(12): p. 1463–1472.

46. Humphrey, W., A. Dalke, and K. Schulten, VMD: visual molecular dynamics. J Mol Graph, 1996. 14(1): p. 33-8, 27-8.

47. Grace, T., GRACE: GRaphing, Advanced Computation and Exploration of data.

48. Inkscape, *Project Inkscape*.

