## Supplementary information for "The evolution and mechanistic versatility of the bacterial NADH dehydrogenases type II"

**Table S1. List from C - COG1252 - NADH dehydrogenase, FAD-containing subunit**

| **Clade** | **Taxonomy Categories** | | **Available in COG** | **Collected initial** | **Clean redundancy (>99% Id)** | **Length seqs** |
| --- | --- | --- | --- | --- | --- | --- |
| Gracilicutes | **Acidobacteria** | | 7/7 organisms 12 genes | 12 | 12 | 263-465 |
| Terrabacteria | **Actinobacteria** | | 137/155 organisms 356 genes | 190 | 190 | 343-649 |
| Gracilicutes | **Aquificae** | | 7/9 organisms 15 genes | 8 | 8 | 388-476 |
| Gracilicutes | **Bacteroidetes** | | 86/107 organisms 132 genes | 165 | 106 | 388-485 |
| Gracilicutes | **Chlamydiae** | | 3/6 organisms 5 genes | 5 | 5 | 209-533 |
| Gracilicutes | **Chlorobi** | | 5/5 organisms 14 genes | 8 | 8 | 399-484 |
| Terrabacteria | **Chloroflexi** | | 8/14 organisms 13 genes | 10 | 10 | 389-468 |
| Terrabacteria | **Cyanobacteria** | | 40/41 organisms 158 genes | 55 | 55 | 373-791 |
| Gracilicutes | **Deferribacteres** | | 5/5 organisms 6 genes | 6 | 6 | 410-458 |
| DST | **Deinococcus-Thermus** | | 6/6 organisms 8 genes | 6 | 6 | 214-465 |
| Terrabacteria | **Firmicutes** | **Bacilli** | 66/73 organisms 142 genes | 96 | 95 | 350-646 |
|  |  | **Clostridia** | 27/79 organisms 37 genes | 30 | 30 | 331-682 |
|  |  | **Negativicutes** | 4/10 organisms 5 genes | 5 | 5 | 418-432 |
|  |  | **Tissierellia** | 2/9 organisms 2 genes | 2 | 2 | 341-355 |
| Fusobacteria | **Fusobacteria** | | 0/6 organisms 0 genes |  |  |  |
| Terrabacteria | **Mollicutes** | | 0/14 organisms 0 genes |  |  |  |
| Gracilicutes | **Planctomycetes** | | 10/14 organisms 14 genes | 9 | 9 | 402-760 |
| Gracilicutes | **Proteobacteria** | **α** | 117/158 organisms 185 genes | 127 | 127 | 348-736 |
|  |  | **β** | 74/102 organisms 123 genes | 62 | 62 | 363-765 |
|  |  | **δ** | 30/39 organisms 51 genes | 37 | 37 | 367-667 |
|  |  | **ε** | 9/12 organisms 32 genes | 8 | 8 | 389-489 |
|  |  | **γ** | 188/224 organisms 296 genes | 190 | 187 | 226-784 |
| Gracilicutes | **Spirochaetes** | | 6/11 organisms 8 genes | 5 | 5 | 395-703 |
| DST | **Synergistetes** | | 0/5 organisms 0 genes |  |  |  |
| DST | **Thermotogae** | | 0/9 organisms 0 genes |  |  |  |
| Gracilicutes | **Verrucomicrobia** | | 6/9 organisms 10 genes | 6 | 6 | 405-523 |
|  | **Other bacteria** | | 14/48 organisms 26 genes | 13 | 13 | 386-477 |

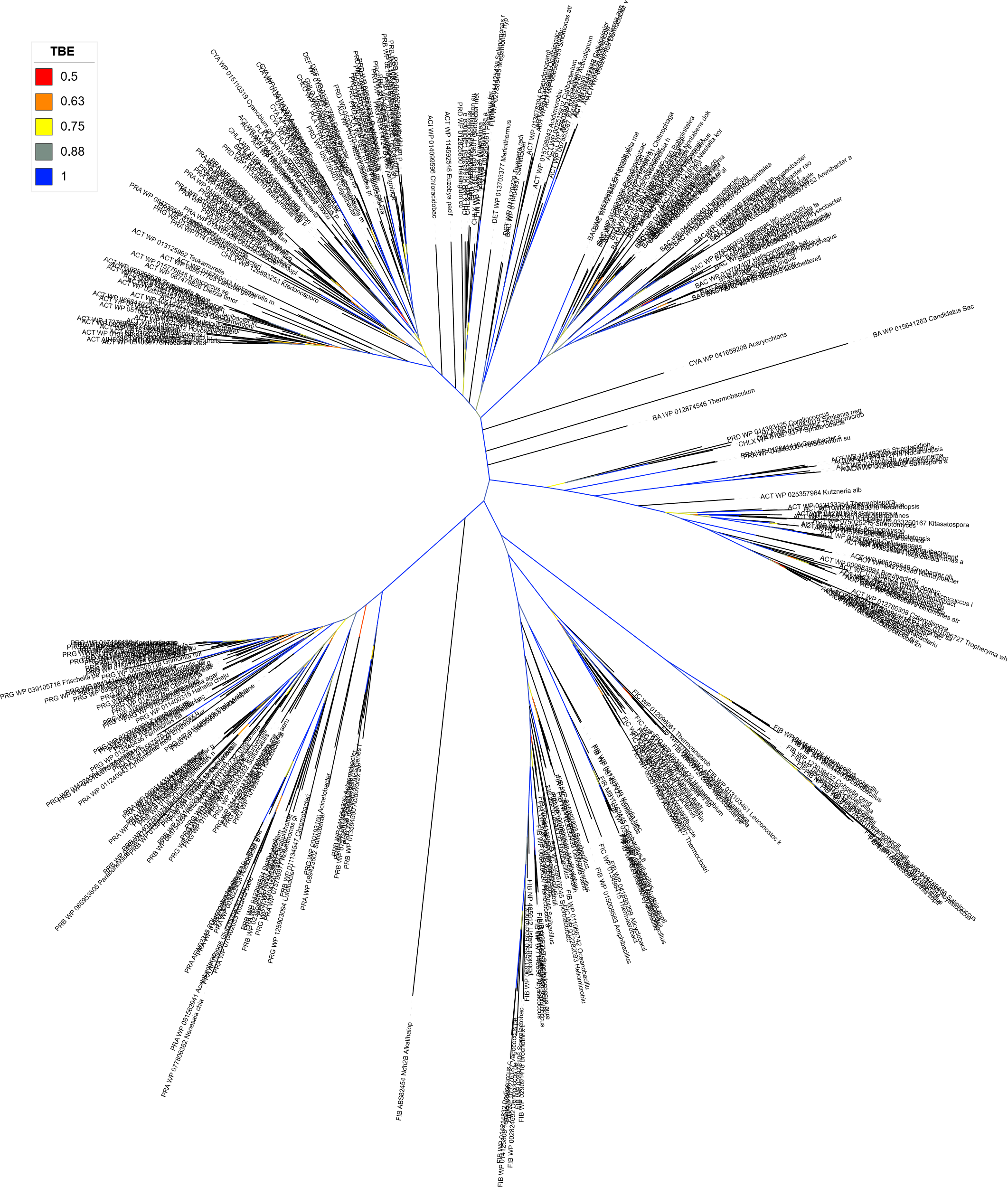

**Fig S1. Fully annotated phylogeny of the bacterial NDH-2 family.** Unrooted tree was obtained in RaxML (MSA 493 sequences, 402 sites). Branches show the TBE values in colors as indicated in the legend. Accession codes and species names are given for each taxa. A tag indicating the association to each bacteria class is given in capitals at the beginning of OTU name as follows: ACI (acidobacteria), ACT (actinobacteria), BAC (bacteroidetes), CHLA (chlamydiae), CHLO (chlorobi), CHLX (chloroflexi), CYA (cyanobacteria), DEF (deferribacteres), DET (deinoccocus-thermus), FIB (firmicutes bacilli), FIC (clostridiales), FIN (negativicutes), PLA (planctomycetes), PRA (α proteobacteria), PRB (β proteobacteria), PRD (δ proteobacteria), PRG (γ proteobacteria), BA (other bacteria).

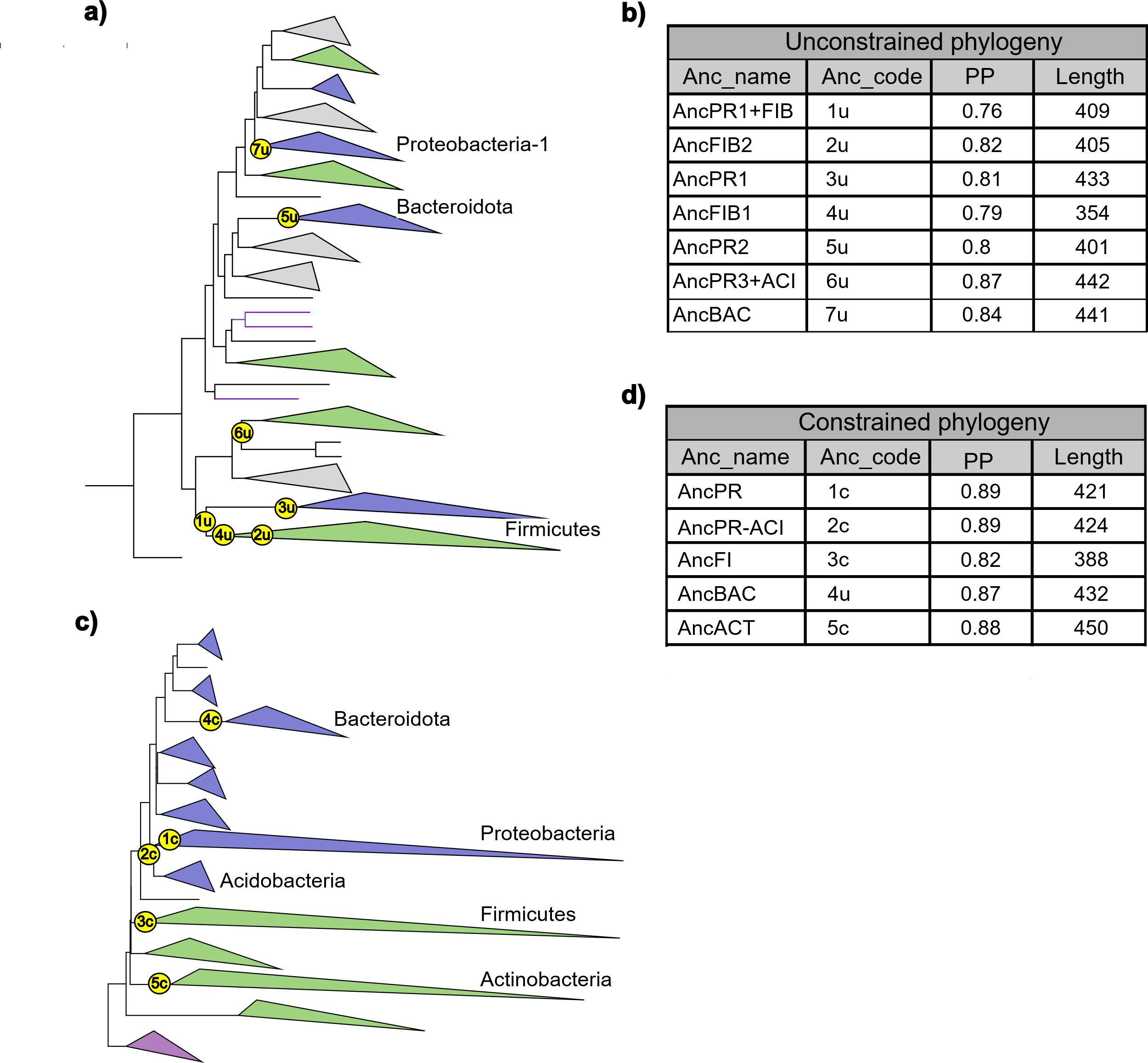

**Fig S2. Ancestral sequence reconstruction. a)** Unconstrained tree topology employed for ASR. Targeted ancestors are shown in yellow circles at the corresponding nodes. Classes in major clades are named, **b)** Summary table for the unconstrained phylogeny targeted ancestors, c) Constrained phylogeny employed in ASR, d) Summary table for the constrained phylogeny targeted ancestors. Color of nodes represent major bacteria clades: gracilicutes (blue), terrabacteria (green), DST (violet). PP: overall posterior probability.

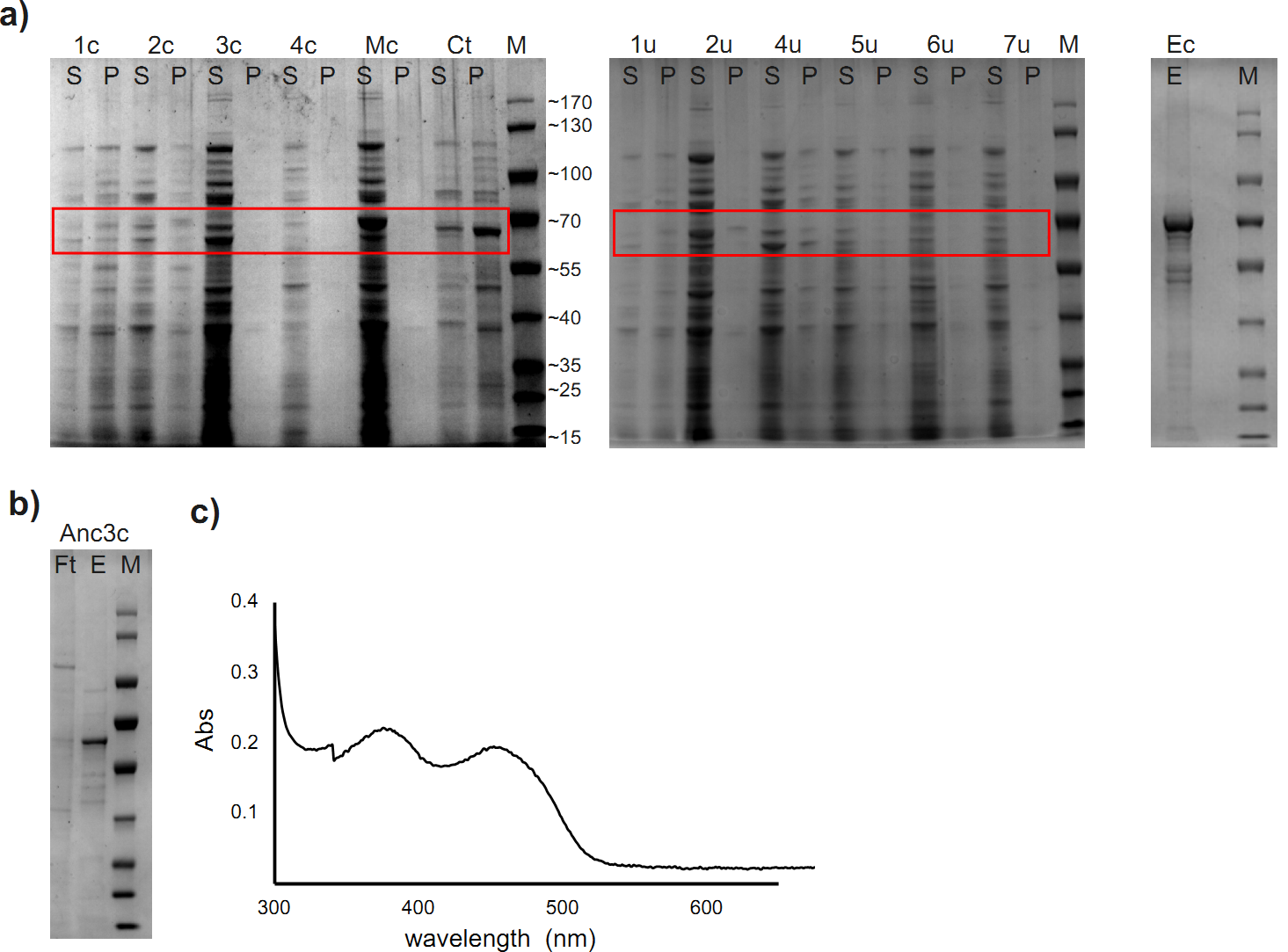

**Fig S3. Enzyme expression and purification. a)** Screening of pBAD constructs expression in 50 mL. S: soluble, P: pellet, M: PageRuler^TM^ prestained protein ladder (Thermo); **b)** purification of Anc3c from 2L culture. Ft: flowthrough, E: elution; **c)** UV-Vis spectra of purified Anc3c.

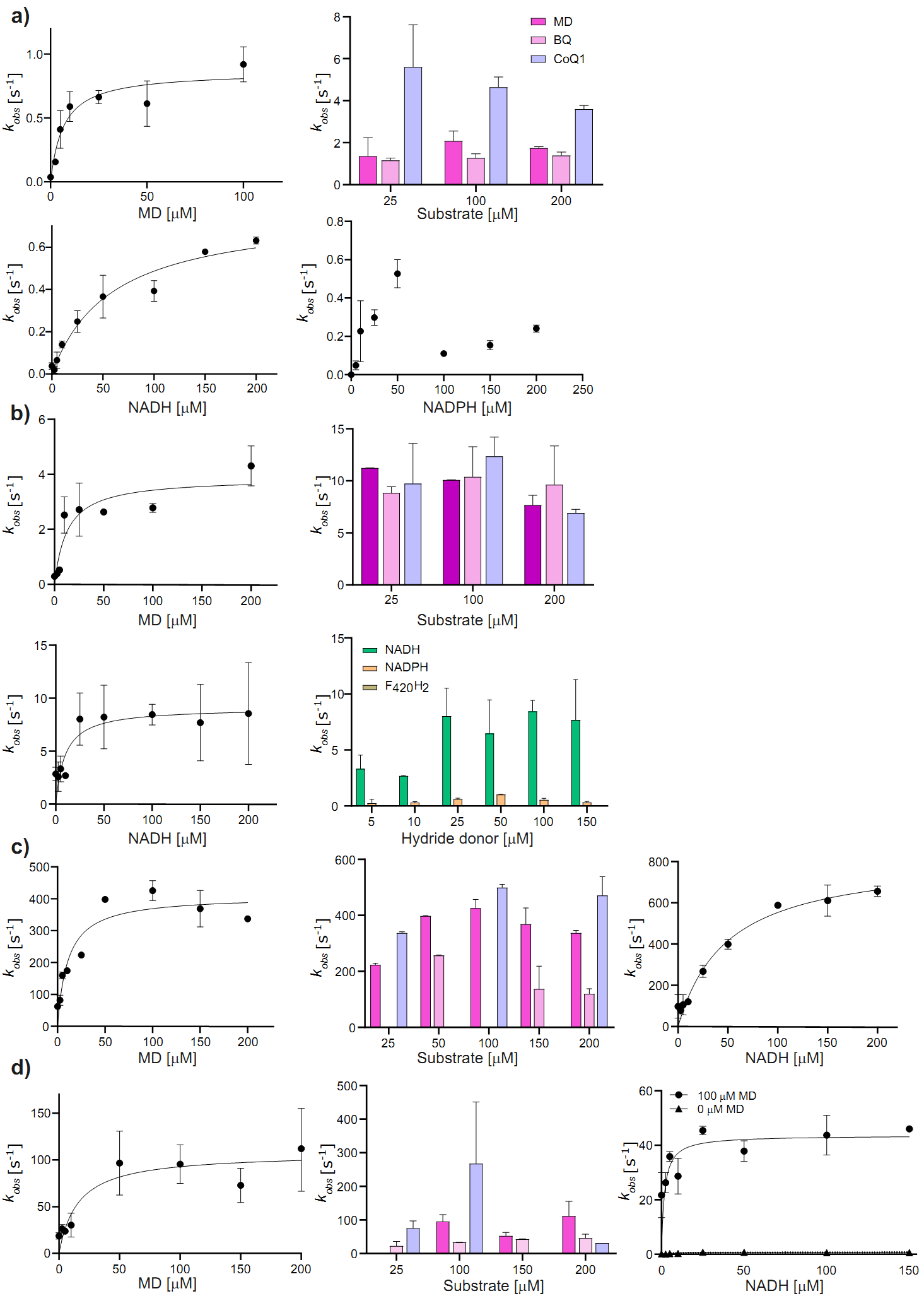

**Fig S4. Steady-state kinetics. a)** Experiments for Anc3c, 0.58-0.1 µM of enzyme were used. MD: menadione, BQ: benzoquinone, CoQ1: coenzyme Q1; **b)** Experiments for Mc-NDH2, 10-100 nM of enzyme were used; **c)** Experiments for Ct-NDH2, 1 nm of enzyme was used; **d)** Experiments for Ec-NDH2, 1 nm of enzyme was used. All experiments were done in triplicates. Mean values are shown as dots and SD values as error bars.

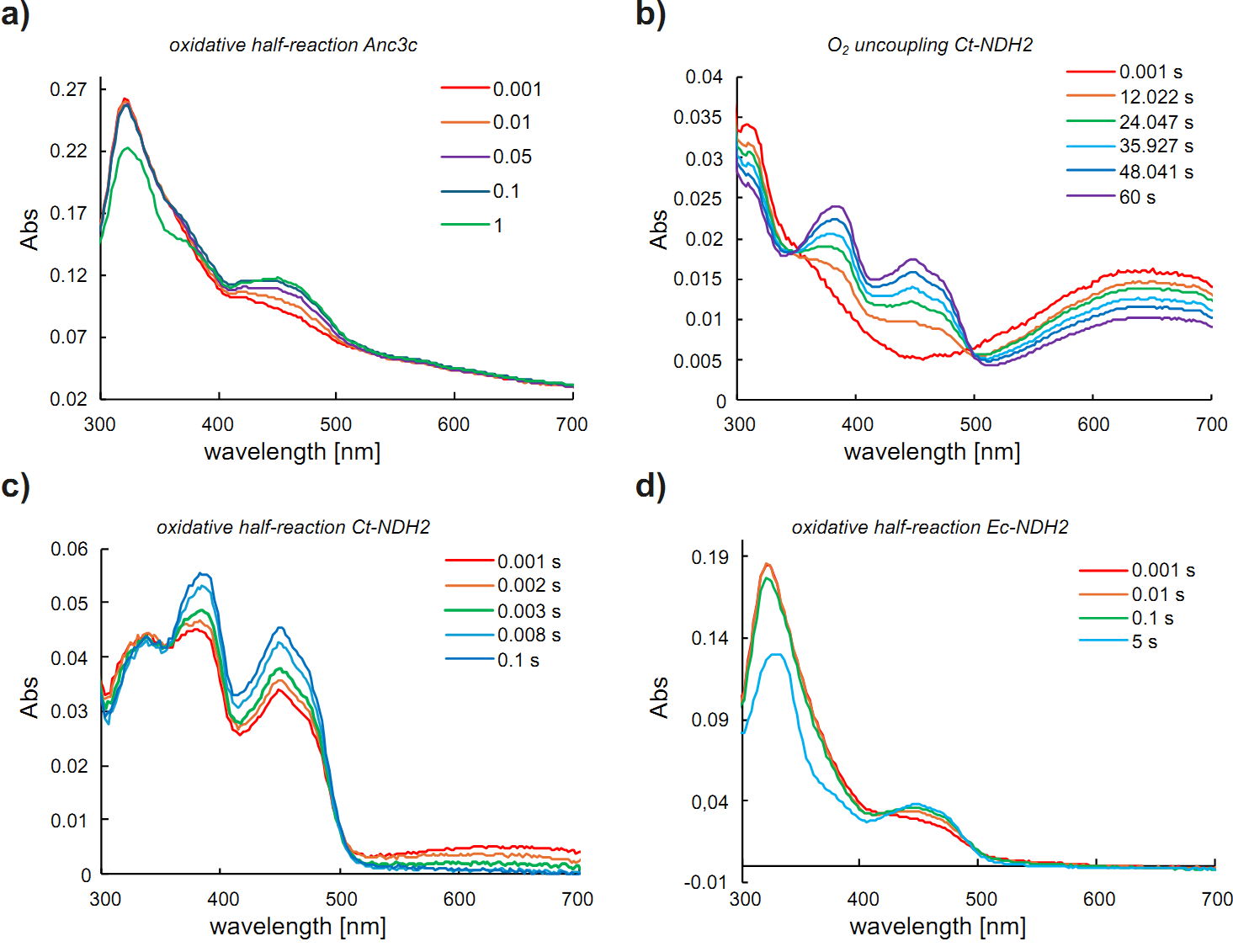

**Fig S5. Pre-steady-state kinetics. a)** Oxidative half-reaction of Anc3c in the presence of menadione; **b)** Oxygen uncoupling reaction of Ct-NDH2 in the absence of substrate; **c)** Oxidative half-reaction of Ct-NDH2 in the presence of menadione; **d)** Oxidative half-reaction of Ec-NDH2 in the presence of menadione. Selected traces are shown. Experiments were repeated three times.

**Table S2. List of new extant NDH-2 selected for experimental characterization**

| **Name** | **Species** | **Accession code** | **Length (aa)** |
| --- | --- | --- | --- |
| AF_NDH2 | *Anoxybacter fermentans* | WP_127015928 | 581 |
| TO_NDH2 | *Thermosediminibacter oceani* | WP_013276488 | 584 |
| BS_NDH2 | *Bacillus subtilis* | NP_391100 | 355 |
| TF_NDH2 | *Thermobifida fusca* | WP_011290879 | 458 |
| MR_NDH2 | *Muricauda_ruestringensis* | WP_041801494 | 435 |
| XO_NDH2 | *Xanthomonas_oryzae* | WP_027704255 | 430 |
| MH_NDH2 | *Marinithermus_hydrothermalis* | WP_013703377 | 430 |

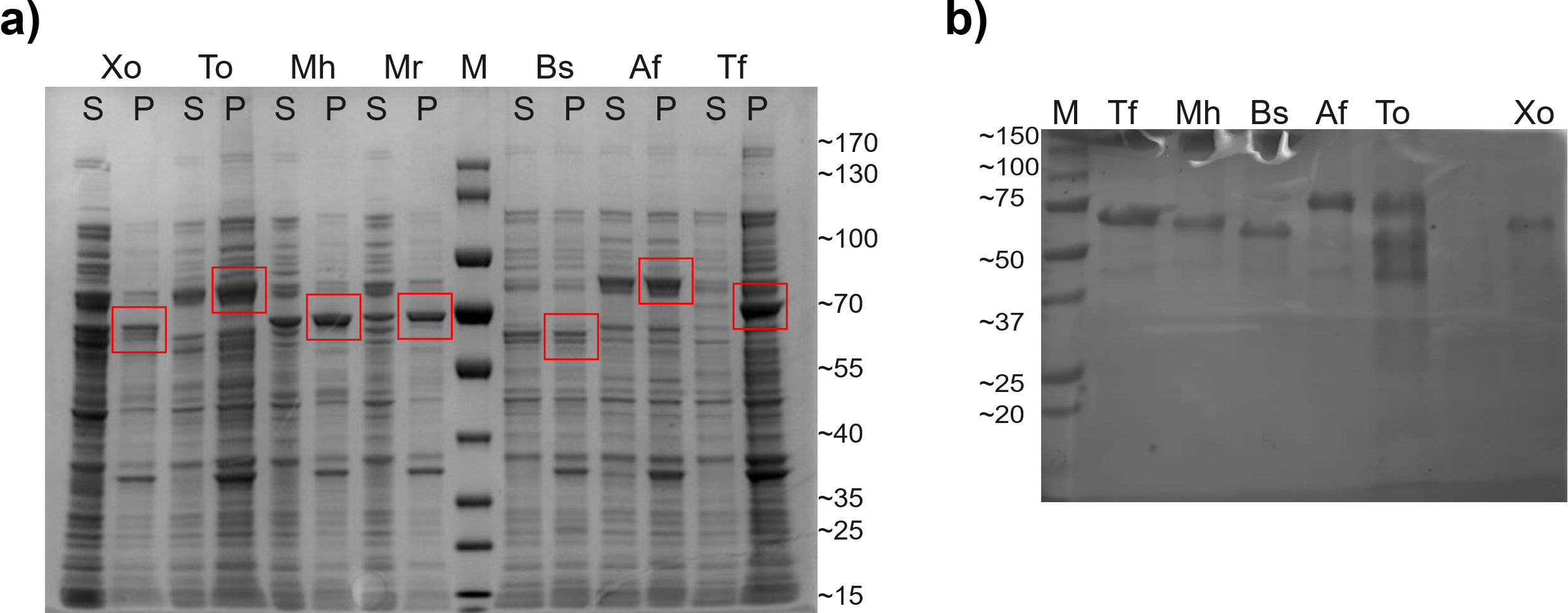

**Fig S6. Expression screening and purification of new extant NDH-2. a)** Screening of pBAD constructs expression in 20 mL. S: soluble, P: pellet, M: PageRuler^TM^ prestained protein ladder (Thermo); **b)** purified proteins by affinity chromatography. M: Precision Plus Protein^TM^ Standards (Bio-rad).

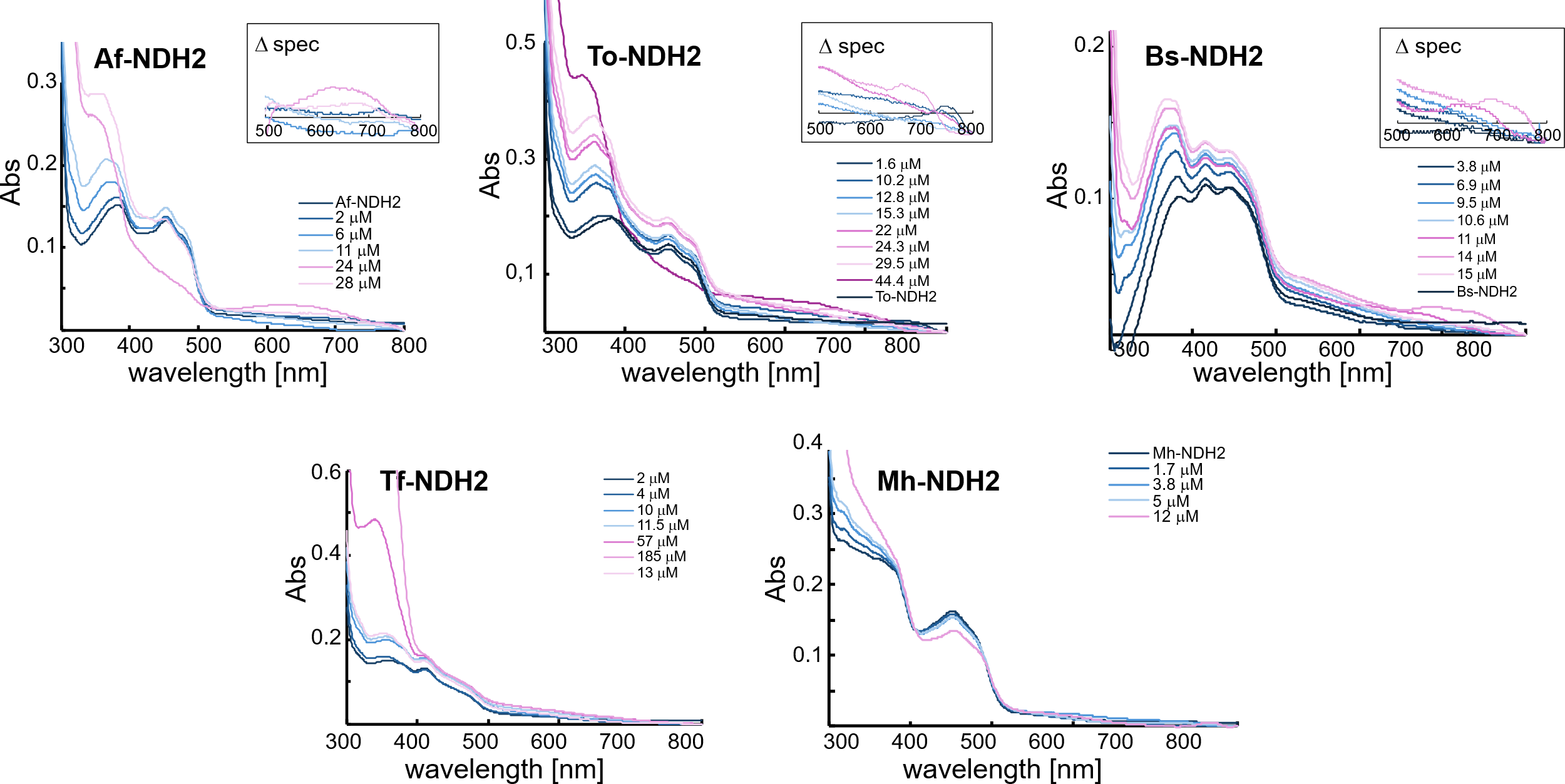

**Fig S7. NADH titration experiments of newly characterized extant NDH-2**. Top row shows enzymes from firmicutes class, forming the CTC upon mixing with 2x molar concentration of NADH. In the insets the difference of spectra among NADH concentrations in the 600-800 nm region is shown. Bottom row corresponds to enzymes from actinobacteria and DET groups, not forming the CTC. Enzyme concentration varied from 6-15 µM.

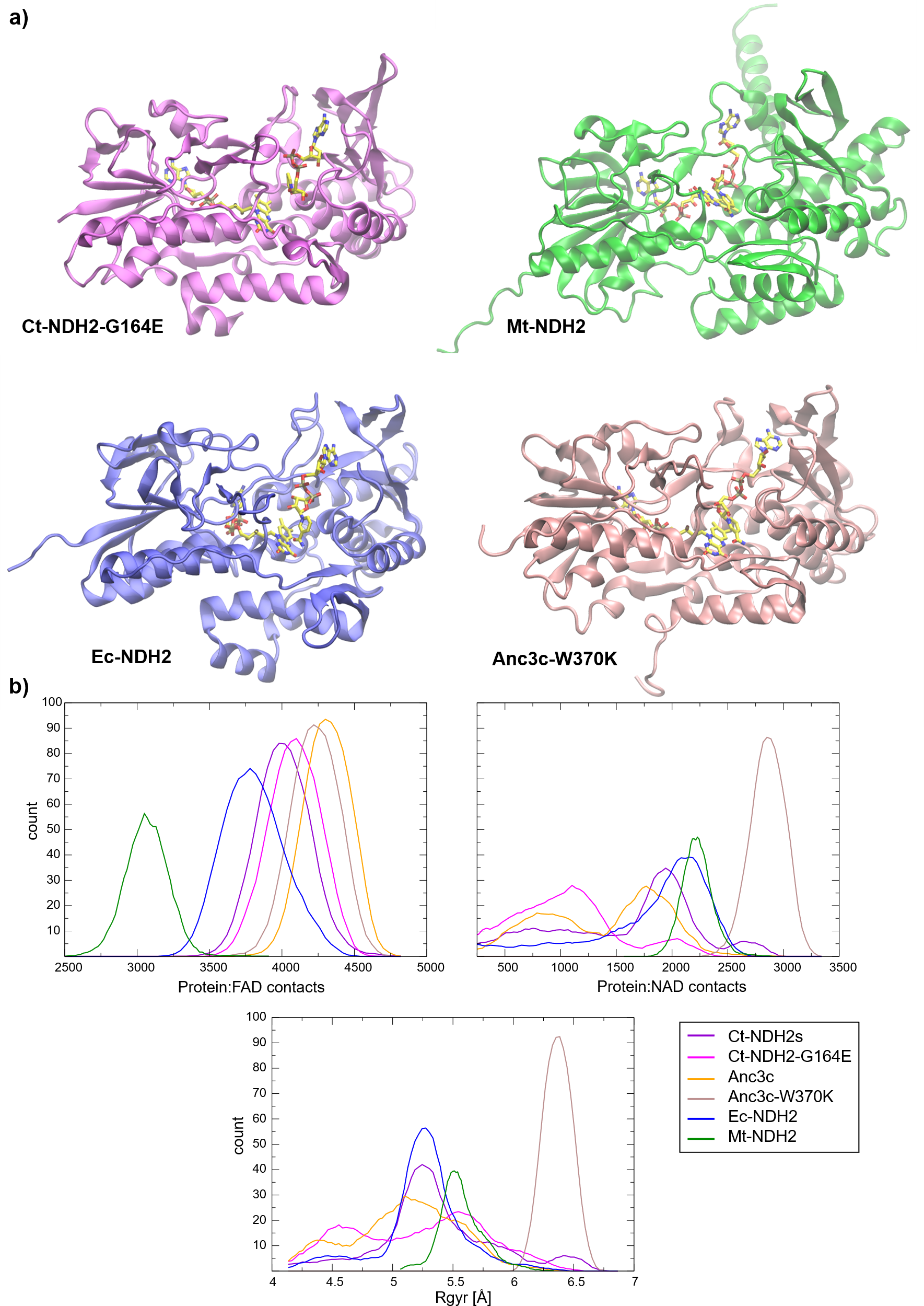

**Fig S8. Selected structures and MD sanity check. a)** Representative structures of Ct-NDH2-G164E (PDB: 5KMP), Mt-NDH2 (Uniprot: P95160), Ec-NDH2 (Uniprot: P00393) and Anc3c-W370K (AlphaFold model) are shown in ribbons and FAD/NAD cofactors in bond representations, with atoms colored as follows: C in yellow, O in red and N in blue. **b)** Graphs show counts of protein:FAD, protein:NADH and the NADH radius of gyration (Å) for all structures investigated across the entire simulation time.

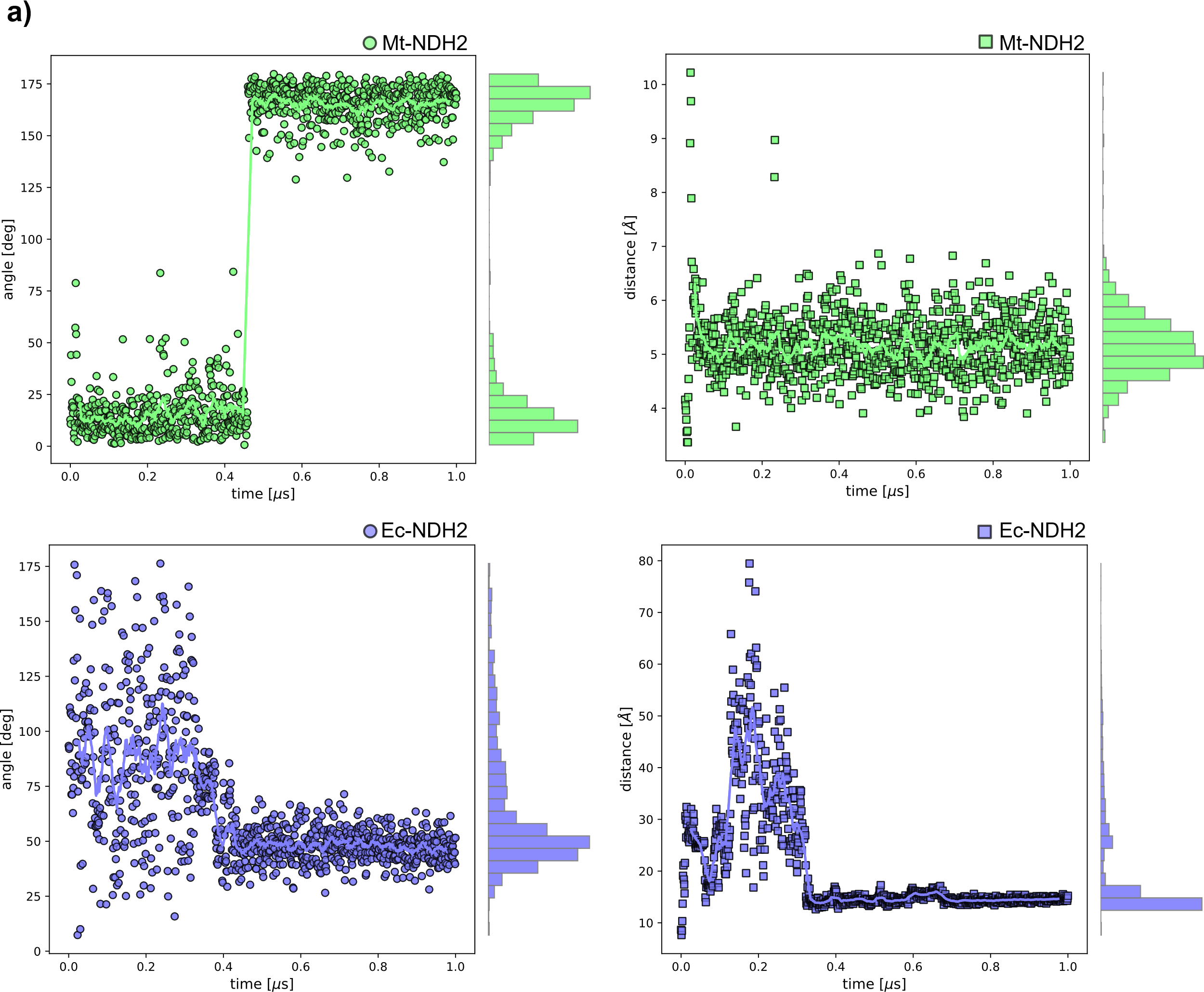

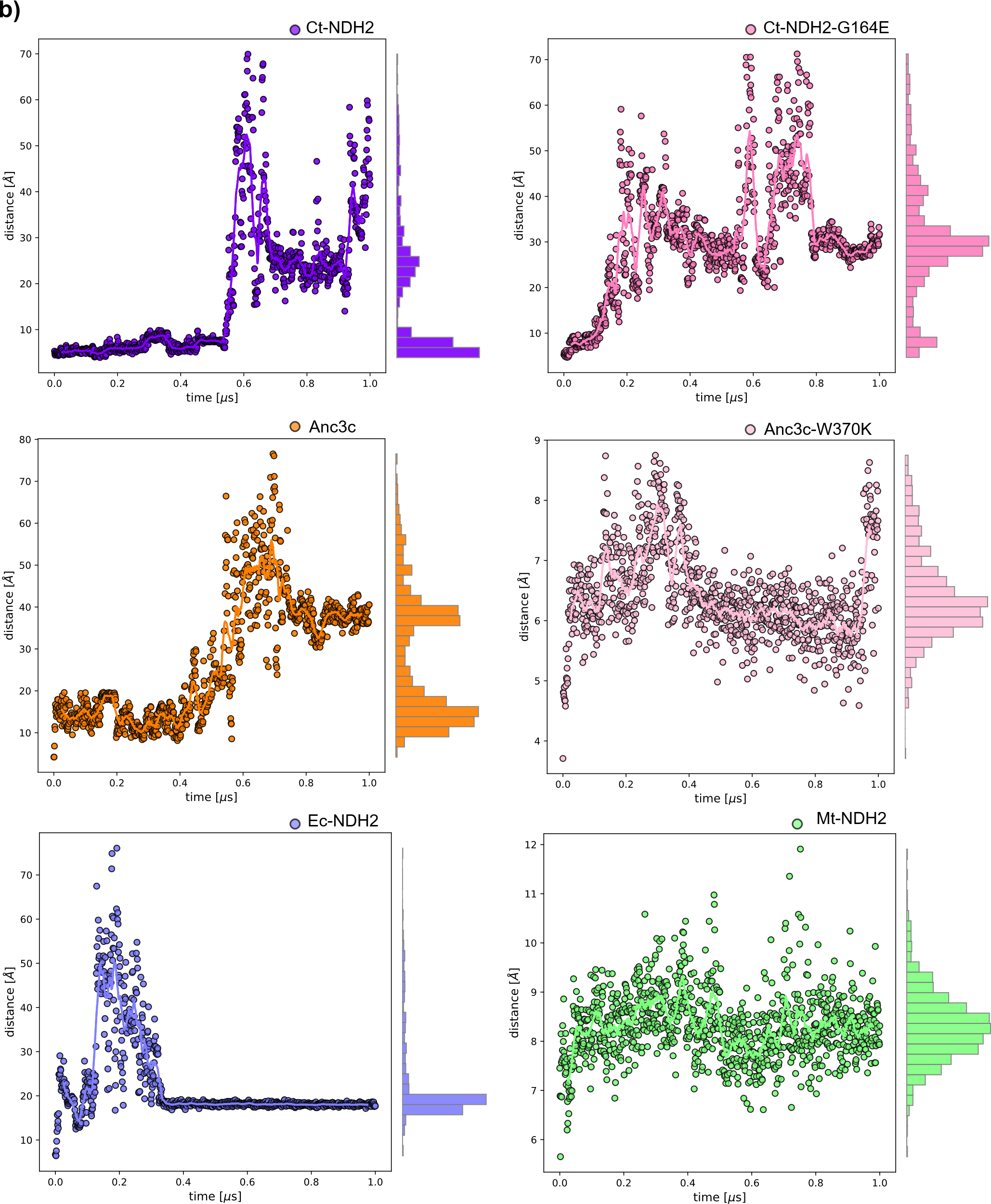

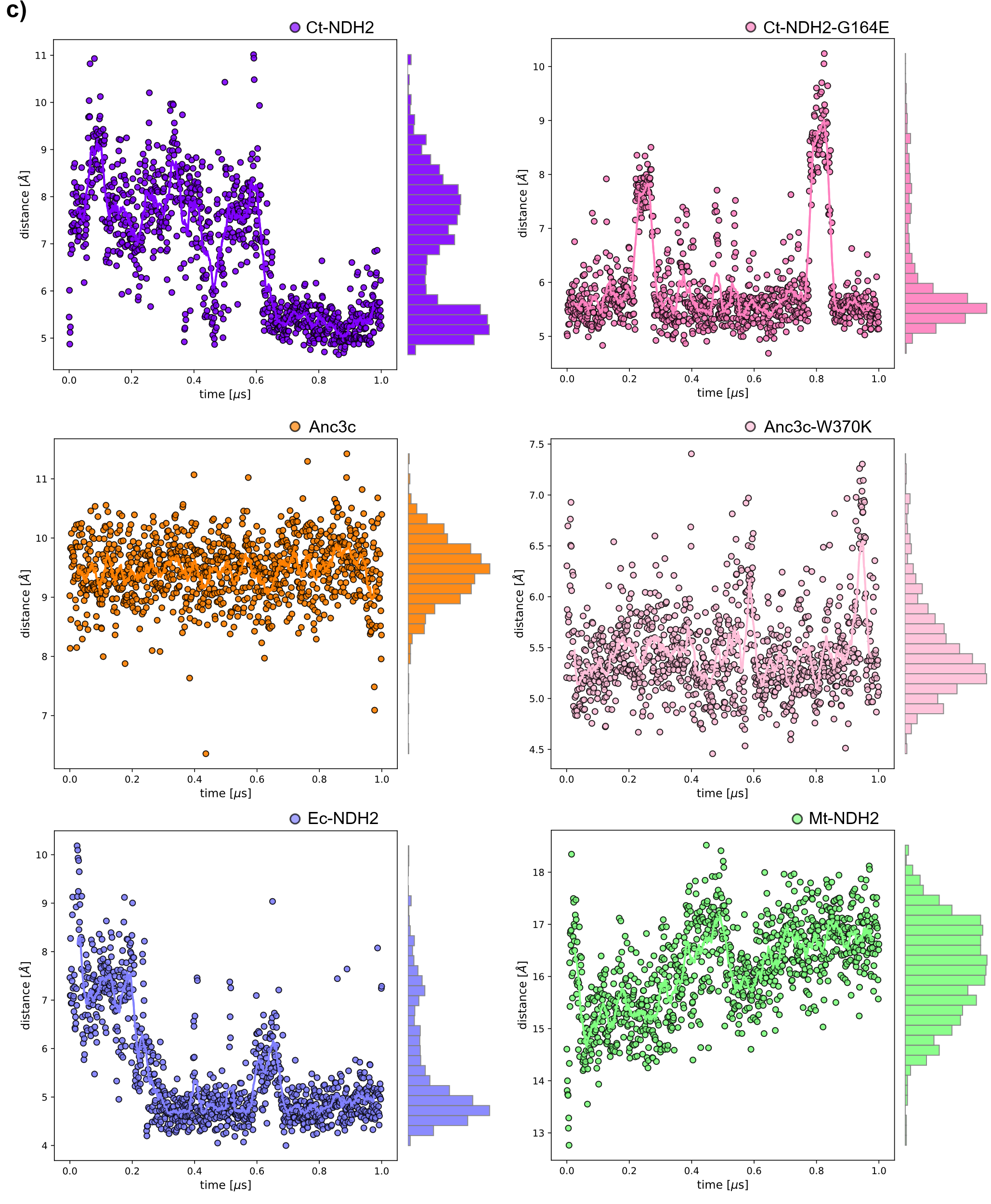

**Fig S9. Selected structures and MD sanity check. a)** Left: angle measurements between the planes formed by the aromatic systems of FAD and NADH in Mt-NDH2 (P95160, in green) and Ec-NDH2 (P00393, in violet) over 1 µs simulation time. Right: minimum distance measurements among the N8 atom from FAD and C19 atom from NADH in Mt-NDH2 and Ec-NDH2 over 1 µs simulation time; **b)** Minimum distance among selected residues to atom N6 of NADH molecule: residue F165 (atom name Cd2) for Ct-NDH2 & Ct-NDH2-G164E, P163 (atom Cg) for Anc3c & Anc3c-W370K, A180 (atom Cb) for Ec-NDH2 and P180 (atom Cg) for Mt-NDH2; **c)** Minimum distance among selected residues to atom N8 of FAD molecule: residue K376 (atom name Nz) for Ct-NDH2 & Ct-NDH2-G164E, W370 (atom Cz2) for Anc3c, K370 (atom Nz) for Anc3c-W370K, Y379 (atom Cz) for Ec-NDH2 and W396 (atom Cz2) for Mt-NDH2. In all cases molecule numbering corresponds to the PDB file.
